# A hypothalamic node for the cyclical control of female sexual rejection

**DOI:** 10.1101/2023.01.30.526259

**Authors:** Nicolas Gutierrez-Castellanos, Basma Fatima Anwar Husain, Inês C. Dias, Kensaku Nomoto, Margarida A. Duarte, Liliana Ferreira, Bertrand Lacoste, Susana Q. Lima

## Abstract

Internal state-dependent behavioral responses, such as the switch between rejection and acceptance of sexual advances depending on a female’s reproductive capacity, are fundamental to maintain social interactions and wellbeing. Here we characterize a dedicated circuit for the cyclical control of rejection behavior in the mouse, located in the anterior ventrolateral ventromedial hypothalamus (aVMHvl). *In vivo* recordings reveal that progesterone receptor expressing neurons of the aVMHvl (aVMHvl^Pr+^) are active during rejection but silent when females accept the male. Moreover, we show that aVMHvl^Pr+^ neurons receive reduced excitatory to inhibitory synaptic input balance during the receptive phase of the reproductive cycle. Finally, optogenetic activation of aVMHvl^Pr+^ neurons in receptive females is sufficient to increase rejection behavior, disrupting the probability of copulation, without affecting other male-directed interactions. This population of aVMHvlPr+ neurons is thus a key neural substrate controlling female sexual behavior, providing an additional barrier to mating when fertilization is not possible.

## Introduction

Cyclic fluctuations in the levels of sex hormones closely coordinate female sexual receptivity with reproductive capacity. This is clearly demonstrated in rodents, as females will only accept male copulation attempts with immobility and lordosis, the sexually receptive posture, and have sex during the fertile phase of the reproductive cycle (physiological estrus or receptive phase). In marked contrast, outside the fertile phase (physiological diestrus or non-receptive phase), copulation never occurs, not only because the presence of a male fails to activate the neural circuits promoting lordosis^1–4^, but also because females actively reject copulation attempts ^5, 6^. Unfortunately, while a lot of attention has been dedicated to the study of female receptivity/lordosis^2, 7–11^, rejection behavior has largely been ignored and often regarded as a mere lack of receptivity.

From a neural perspective, it is long established that the ventrolateral region of the ventromedial hypothalamus (VMHvl) holds a crucial role in the control of female sexual receptivity: besides receiving male-derived information, such as olfactory signals^2, 12–14^ or the flank somatosensory stimulation derived from copulation attempts^15, 16^, electrical stimulation of the VMHvl increases lordosis display while its electrolytic lesion has the opposite effect^7, 8^. In addition, the expression of receptors for female sex hormones estrogen (E) and progesterone (P) is observed in a vast percentage of VMHvl neurons (ER+ and PR+ neurons)^11, 12^. ER+ and PR+ VMHvl neural populations are necessary for lordosis expression^11, 12^ and their activation through local infusion of sex hormones in the VMHvl is sufficient to restore sexual receptivity in ovariectomized rodents^17^. Sex hormones affect neuronal physiology in various ways^4, 5^, but given that E and P levels continuously change across the cycle and their effects are concentration dependent, neuronal properties are in continuous flux as well. In fact, several studies show that the electrophysiological and structural properties of VMHvl neurons are modulated by E and P and vary across the cycle^2, 14, 18–21^. These results strongly suggest that the computations performed by VMHvl neurons may change dynamically across the reproductive cycle, biasing the outcome of socio-sexual interactions in coordination with female reproductive capacity. Indeed, recent evidence links the cyclical nature of sex hormone concentration with VMHvl neuronal properties and behavior, revealing that fertile levels of E and P are associated with synaptic input potentiation and structural potentiation of the output of defined neural subpopulations crucial for the expression of sexual receptivity^2, 13^.

While the receptivity dial has been associated with plasticity of VMHvl inputs/outputs, the active rejection displayed by non-sexually receptive females remains short of a neuronal substrate. Still, recent studies have uncovered an unappreciated complexity and multifunctional nature of the VMHvl, with the existence of distinct sub-compartments across its anterior-posterior (AP) axis that exhibit different connectivity^22^, transcriptomic profiles^23^, electrophysiological properties^14, 18^, response to fluctuating levels of sex hormones^15^ and function^24^. Accordingly, recent evidence suggests that the control of sexual receptivity, once thought to involve the entire VMHvl, is instead restricted to its most posterior region^2, 13^. Even more surprising was the recognition that maternal aggression is also controlled by the posterior VMHvl, in a hormonal state–dependent manner^12, 25^, prompting us to speculate if sexual rejection could also be controlled by this hypothalamic structure. Sexual rejection, however, should not be mistaken for aggression. In fact, only lactating mothers seem to exhibit natural aggression^12, 25^. In contrast, rejection behavior in the sexual context is observed in response to the male attempts at copulation and is therefore more accurately described as an active defensive strategy to avoid unwanted social interactions. Interestingly, this form of rejection behavior exhibited by female rodents bears a striking resemblance to the behavior displayed by submissive males in the presence of dominant male intruders, including an upright posture, escaping, dashing, boxing and kicking^24, 26^. Importantly, in males, self-defense towards conspecifics is controlled by the anterior portion of the VMHvl ^24^, and as such we asked if anterior neurons in the female VMHvl could also be involved in rejection behavior when non-receptive.

Here we propose a new role for anterior VMHvl PR+ neurons (aVMHvl^Pr+^) in the cyclical control of female rejection behavior. Evidence supporting this hypothesis first came from the observation that the aVMHvl is more active when non-receptive females interact with a male, an interaction dominated by rejection behavior. Taking advantage of calcium imaging methods to monitor neural activity across the reproductive cycle, we found spatial segregation in the response profile of VMHvl^Pr+^ neurons across the AP axis: while lordosis-related activity seems to be localized to PR+ cells in the posterior VMHvl (pVMHvl^Pr+^), corroborating existing literature, the aVMHvl^Pr+^ population is active when non-receptive females reject the male’s copulation attempts, but remains silent when a receptive female is mounted by the male. Electrophysiological characterization of the properties of PR+ neurons across the reproductive cycle revealed periodic and spatially segregated changes in their incoming synaptic input. In particular, aVMHvl^Pr+^ neurons seem to experience profound changes in the amount of excitatory and inhibitory input they receive, such that the excitation/inhibition (E/I) ratio of these cells is markedly higher in non-receptive females than in receptive females. This change in E/I balance suggests that aVMHvl^Pr+^ neurons might be more active when a non-receptive female interacts with a male than when a receptive female does, corroborating our initial functional efforts. To causally relate these findings, we performed localized optogenetic stimulation of aVMHvl^Pr+^ neurons in receptive females, which normally exhibit very low levels of rejection. As expected, our manipulation led to increased rejection behavior in receptive females, fully preventing mating in some couples, while significantly delaying the progression to sexual consummation in others, establishing a role for the aVMHvl^Pr+^ population in the control of rejection behavior. This leads us to propose that the female’s response to copulation attempts across the reproductive cycle is the outcome of two parallel processes, receptivity and rejection, both controlled by distinct and spatially segregated VMHvl^Pr+^ populations, whose activity is modulated by sex hormones in a bidirectional and opposing manner.

## Results

### PR+ neurons in the anterior, but not posterior, VMHvl are preferentially active when diestrus females reject copulation attempts from the male

To explore a possible role of the VMHvl in the control of rejection behavior, we began by comparing the activity elicited in the VMHvl of two separate groups of females at polar opposites of the reproductive cycle after an interaction with a male. Once opposite sex mice get together, and after an initial period of social investigation (which in laboratory mice and in non-naturalistic conditions seems largely unaffected by the female’s reproductive state^14, 27^), the male will attempt copulation by placing his front paws on the female’s flanks. The female’s response to copulation attempts however is drastically different across the reproductive cycle, which lasts 4–5 days in mice and is characterized by 4 distinct phases that may be determined according to the cell types observed in a vaginal smear^5^ (Fig. 1A). In naturally-cycling females, outside of the fertile phase of the cycle, copulation attempts are rejected without exception, by walking/running away and the display of a variety of active defensive-like behaviors, including kicking and boxing^5, 6, 14^ (Fig. 1A, Video 1). In contrast, when fertile, a short period of time between the proestrus and estrus phases (PE), females eventually accept the male, allowing penile insertion, several mounts and ejaculation^6^ (Fig. 1A). The first attempt at copulation can therefore be used to identify a transition in the male’s sexual arousal, whereas the moment when the female accepts him sets the start of the classically defined consummatory phase. As the dichotomization of the behavior is bound to two critical steps, first the male’s copulation attempt and second her acceptance, here we divided the social interaction into 3 phases (instead of the classical division into appetitive/consummatory behavior^28, 29^): appetitive 1 (until the first mount attempt, App1), appetitive 2 (until the first successful mount, App2) and consummatory (Cons) (Fig. 1A, see legend for details).

**Figure 1.**
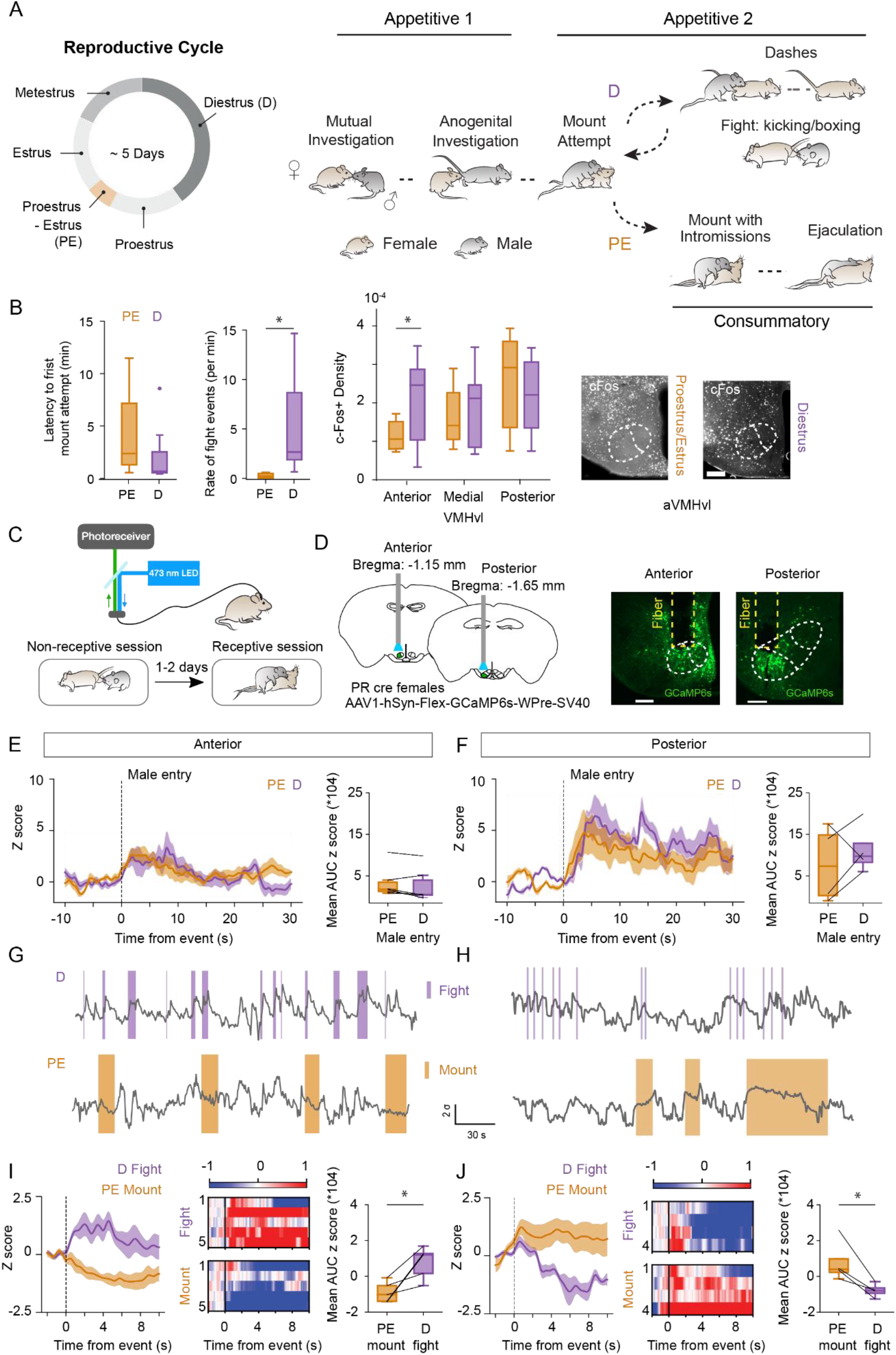
VMHvl neuronal activity during sexual acceptance and fighting behavior along the antero-posterior axis in different internal states. (A) Left: Schematic diagram of the female reproductive cycle indicating, in orange, the transition between proestrus and estrus which corresponds to the period of peak female fertility and sexual receptivity. Right: Simplified diagram of the structure of sexual behavior and its division into 3 different phases — 1. appetitive 1 (App1): this phase starts at the entry of the male into the experimental cage, includes the initial investigatory period between the male and female, and concludes as the male performs his first mount attempt; 2. appetitive 2 (App2): this phase starts at the first failed mount attempt by the male and continues until the first successful mount with thrusts. Whenever the first mount attempt leads to a successful mount with thrusts, this phase is absent; 3. consummatory (Cons): this phase starts at the beginning of the first mount with thrusts and ends when the male ejaculates. D females do not have Cons considering there is no occurrence of a mount with thrusts. (B) Left: Latency to first mount attempt and fighting rate in the behavioral sessions followed by cFos detection. Right: cFos-positive cell density in the VMHvl along the anterior-posterior axis in either PE (orange) or D (purple) females after mounting or fighting behavior, respectively, and representative images showing cFos expression in the anterior VMHvl in each condition. Scale bar: 200 µm. (C) Schematic illustrating photometry recordings and experimental design. (D) Left: Viral construct and implantation coordinates in PR Cre females in either the anterior (n = 6) or posterior (n = 8) VMHvl. Right: Representative images showing GCaMP6s expression and fiber placement in anterior and posterior VMHvl. Scale bars: 200 µm. (E,F) Left: averaged peri-event time histograms (PETHs) of z-scored GCaMP6s data from anterior (F) or posterior (G) recordings aligned to male entry during the D and PE session (mean ± sem). Right: comparison of area under the curve of the z score within 30 seconds after male entry between D and PE sessions. D: purple, PE: orange. (G,H) Representative z-scored normalized GCaMP6s fluorescence traces (gray) from PR Cre females implanted in the anterior (G) or posterior (H) VMHvl during fights (purple shaded region) and mounts (orange shaded region) in D and PE sessions, respectively. (I,J) Left: averaged PETHs of z-scored GCaMP6s data from anterior (I) or posterior (J) recordings aligned to male entry, fights or mounts during the D and PE session (mean ± SEM). Middle: heatmaps of averaged PETHs aligned to various behaviors. Right: comparison of area under the curve of the z score within 5 seconds after the event between D and PE sessions. D: purple, PE: orange; *p<0.05, paired t-test.

To explore the activation of VMHvl neurons during opposite phases of the cycle we used diestrus females (D) as non-fertile/non-receptive and PE as fertile/receptive females and the immediate early gene cFos as a proxy for neural activity^30^. Despite the fact that males attempted copulation with similar latencies in both groups of females (Fig. 1B), kicking and boxing (which here we denote as fighting behavior) was 21 fold more frequent in D than in PE females (Fig. 1B, PE rejection rate: median [IQR]=0.44 [0.17 0.55] events per minute, D rejection rate: median [IQR]=2.59 [1.82 8.55] events per minute; Mann–Whitney U test, p=0.0004; D/PE (mean ratio)=21.22); and while all PE females had sex, D females actively rejected the male without exception until the session was interrupted, deterring any possibility of copulation. The opposing behavioral profiles observed in PE versus D females was accompanied by a distinct pattern of VMHvl activation at its most anterior subdivision (Fig. 1B; 2-way ANOVA, reproductive phase x AP level interaction factor, F(2,16)=4.22, p=0.03). Specifically, a significantly higher number of aVMHvl neurons were active in D females interacting with and rejecting the male when compared to the number of active neurons in the aVMHvl of PE females that mated (aVMHvl cFos density for PE: median [IQR]=1*10^−4^ [0.8*10^−4^ 1.5*10^−4^] cFos+ per mm^2^, and for D=2*10^−4^ [1*10^−4^ 2.9*10^−4^] cFos+ per mm^2^; Sidak’s multiple comparison test, p=0.02). In addition, no difference was observed across the two conditions in the dorsomedial portion of the VMH throughout its AP extension (Fig. S1). Interestingly, in a separate experiment we observed that non-primed ovariectomized (OVX) females, the routinely used model system in studies of female sexual receptivity^31, 32^, exhibit much lower levels of fighting behavior once males attempt copulation when compared with naturally-cycling D females (referred as NC-D in this experiment) (Fig. S2; OVX median [IQR]=0.56 [0.25 0.8] events per minute and NC-D median 1.42 [1.1 1.8] events per minute; unpaired t-test, p=0.02), possibly explaining why the active defensive-like fighting behavior in response to copulation attempts has been largely ignored in the field of female sexual behavior.

Although the cFos experiment revealed that distinct and spatially segregated regions of the VMHvl are activated at different phases of the reproductive cycle, this tool lacks cellular and temporal specificity. Therefore, we next employed the calcium sensor GCaMP6 in a genetically delineated population of VMHvl neurons to monitor the activity of its anterior or posterior subdivisions with fiber photometry while PE or D females interacted with a male (Fig. 1C). We focused our efforts on the progesterone receptor–expressing (PR+) population which is crucial for female sexual behavior^2, 11^. AAV vectors expressing GCaMP6 in a Cre-dependent manner were unilaterally injected either in the anterior or posterior part of the VMHvl of PR-Cre females, and implanted with a fiber optic cannula above the injection location for imaging (Fig. 1D). Once these females recovered from surgery, they were allowed to freely interact with a male during 2 separate sessions — one during the D phase, followed by one during the PE phase of the reproductive cycle (Fig. 1C). As before, D females remained in App2 and the interaction was characterized by a series of failed attempts at copulation by the male and fighting behavior by the female (Fig. S3). On the other hand, for couples where the female was in the PE phase, the interaction progressed through all behavioral epochs, including successful mounts with thrusts during Cons (Fig. S3).

Male chemosignals elicited activation of both aVMHvl^Pr+^ and pVMHvl^Pr+^ neurons, as shown by a marked increase in fluorescence upon male entry in the experimental arena. The magnitude of this response was constant irrespective of the reproductive state (Fig. 1E,F). In contrast, the activity throughout the AP axis of the VMHvl was significantly different during fighting and mounting behaviors which characterize the opposite reproductive states (Fig. 1G-J). While aVMHvl^Pr+^ cells were active when D females fought the male, the activity of these neurons was inhibited when PE females were successfully mounted by the male (Fig. 1G,I; aVMHvl^Pr+^ PE-mount median [IQR]=–1.02 [–1.41 –0.29] arbitrary units/ a.u. ; D-fighting=1.20 [0.17 1.48] a.u.; paired t-test, p=0.02; and Fig. S4). On the other hand, pVMHvl^Pr+^ neurons showed no significant activation when D females fought the male, but showed robust activation during successful mounts, in line with the well-established role of this sub-region in the control of female receptivity (Fig. 1H,J; pVMHvl^Pr+^ PE-mount median [IQR]= 0.39 [–0.01 2.05] a.u. ; D-rejection=–0.79 [–1.15 –0.41] a.u.; paired t-test, p=0.02)^2^. The fluorescence emitted by aVMHvl^Pr+^ neurons remained decreased during the whole duration of the mount, only returning to baseline levels after the mount was terminated, suggesting that the activity of this sub-region is inhibited while the female is mounted by the male (Fig. S4). These results corroborate our hypothesis that aVMHvl^Pr+^ neurons are tuned to sexual rejection behavior and suggest that their activity must be decreased for copulation to occur. Importantly, fighting and mounting contribute to the strongest and sharpest bouts of acceleration and therefore both types of behavior act as an internal validation mechanism, within and across different recordings, to exclude the existence of movement-induced fluorescence changes. In addition, other behaviors which are also accompanied by sharp motion around the head implant, such as head turns, did not elicit any change in GCaMP fluorescence (Fig. S5).

### PR+ neurons in the anterior, but not posterior, VMHvl receive a differential excitation/ inhibition synaptic balance throughout the reproductive cycle

So far, our results show that the response magnitude of aVMHvl^Pr+^ neurons to male sexual solicitations undergoes reproductive state–correlated changes and is associated with fighting behaviors during the D phase. Several mechanisms may underlie the distinct pattern of activation observed in the aVMHvl^Pr+^ population that we observed with cFos and fiber photometry, such as changes in the incoming synaptic input elicited by close interactions with the male, or by a difference in the ability of aVMHvl^Pr+^ neurons to transform the input received into output. Given that aVMHvl^Pr+^ neurons have comparable intrinsic excitability across the reproductive cycle^18^, we sought to monitor the synaptic inputs arriving at aVMHvl^Pr+^ neurons across different phases of the reproductive cycle.

For this, we performed whole-cell voltage-clamp electrophysiological recordings of spontaneous synaptic currents in acute slices obtained from PE (n=22 mice) or D (n=27 mice) adult PR-EYFP female mice (Fig. 2A). By clamping at the reversal potential of GABAa receptors (–70 mV) and AMPA receptors (0 mV) we were able to isolate spontaneous AMPA glutamatergic (excitatory postsynaptic currents, EPSCs) and GABAergic synaptic events (inhibitory postsynaptic currents, IPSCs). Our results reveal a marked shift in the frequency of spontaneous excitatory and inhibitory synaptic input arriving to PR+ neurons, which was particularly pronounced in the aVMHvl (Fig. 2B-F).

**Figure 2.**
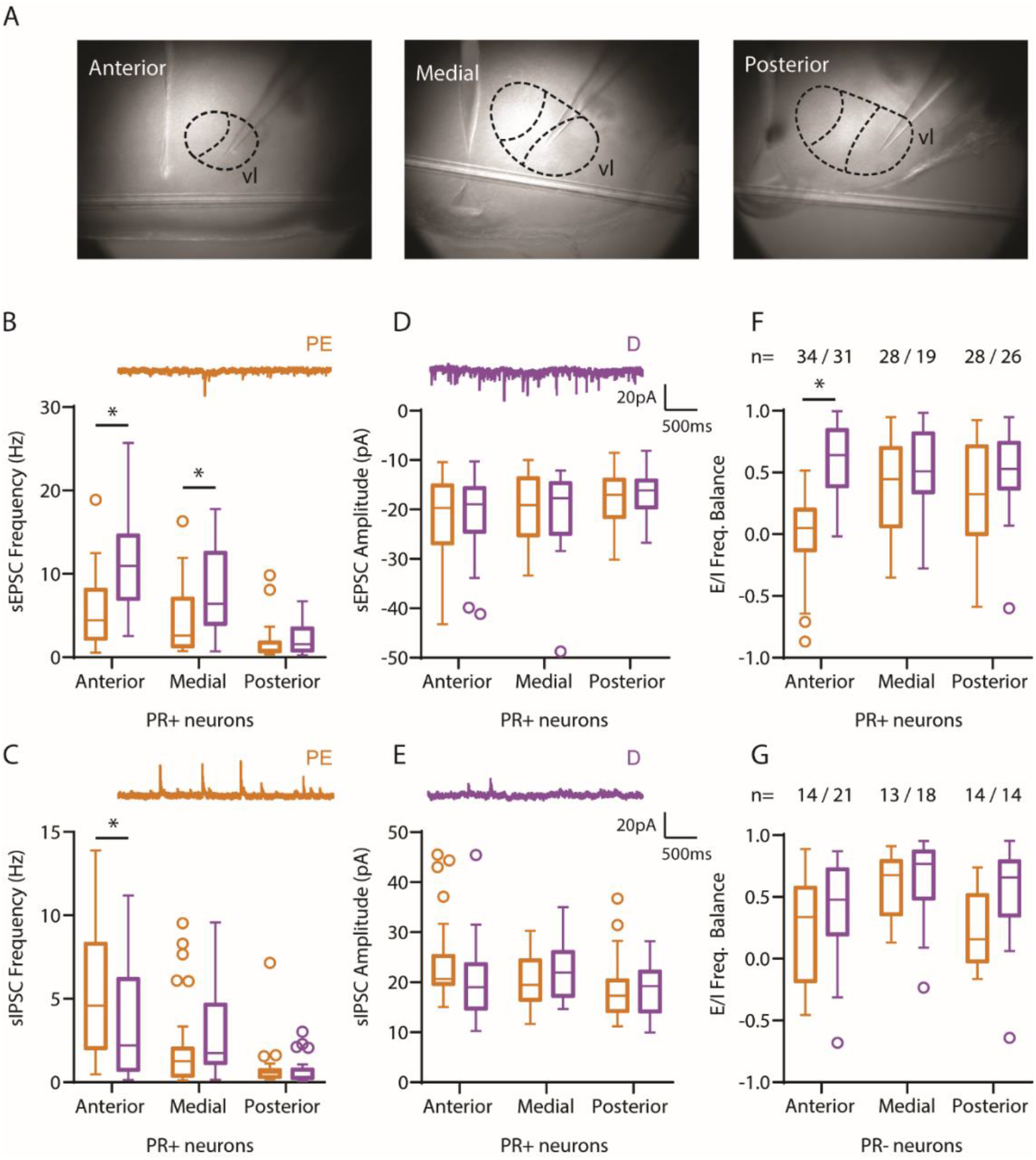
Anterior, but not posterior, VMHvl^Pr+^ neurons undergo synaptic changes across the female reproductive cycle. (A) Pictures of representative anterior, medial and posterior levels of the VMHvl used for electrophysiological recordings. (B-C) Frequency of excitatory and inhibitory spontaneous synaptic events throughout the AP extension of the VMHvl in D and PE females (D-E) Amplitude of excitatory and inhibitory spontaneous synaptic events throughout the AP extension of the VMHvl in D and PE females. (F-G) The synaptic changes observed yielded a change in the excitatory/inhibitory frequency balance of synaptic input of aVMHvl neurons that was specific to PR+ neurons (F), as it was not observed in PR– neurons (G).

When recording the synaptic inputs impinging on VMHvl neurons from PE females, we observed that anterior and medial PR+ neurons receive a significantly lower frequency of EPSCs compared to PR+ neurons from D females (Fig. 2B; aVMHvl^Pr+^ PE median [IQR]=4.43 [2.11 8.21] Hz; D= 10.96 [6.89 14.72] Hz; mVMHvl PE=2.61 [1.22 7.12] Hz; D=6.4 [3.87 12.63] Hz; 2-way ANOVA, reproductive phase factor, F(1,160)=23.5, p<0.0001; Sidak’s multiple comparison test, anterior PE vs. D p<0.0001 and medial PE vs. D p=0.0056). Moreover, anterior, but not medial or posterior, VMHvl^Pr+^ neurons from PE females receive a significantly higher number of spontaneous IPSCs compared to neurons from D females (Fig. 2C; aVMHvl PE median [IQR]= 4.59 [1.99 8.36] Hz; D=2.23 [0.7 6.24] Hz; 2-way ANOVA, reproductive phase x AP interaction factor, F(2,160)=3.71; p=0.02; Sidak’s multiple comparison test, anterior PE vs. D p=0.0052). Changes in the frequency of synaptic input were not accompanied by changes in the amplitude of neither excitatory nor inhibitory synaptic events (Fig. 2D–E). The shift in the frequency of incoming synaptic input yielded a significantly reduced E/I frequency balance in anterior but not medial or posterior, PR+ neurons (Fig. 1F; aVMHvl^Pr+^ PE median [IQR]= 0.05 [–0.13 0.21] a.u.; D=0.64 [0.38 0.85] a.u.; 2-way ANOVA, reproductive phase x AP interaction factor, F(2,160)=7.42; p=0.0008; Sidak’s multiple comparison test, anterior PE vs. D p<0.0001). Changes in E/I balance were specific to PR+ neurons, as they were not present in neurons that did not express EYFP under the PR promoter (PR– neurons, Fig. 2G and Fig. S6A–D).

Changes in the frequency of synaptic currents can result from modifications in presynaptic firing frequency, postsynaptic unsilencing of synapses or changes in presynaptic release probability^33^. To parse out the contribution of these mechanisms, we first recorded miniature synaptic events (mEPSCs and mIPSCs, Fig. S6E–I) in the presence of the sodium channel blocker tetrodotoxin (TTX, 1 μM) to test whether changes in presynaptic firing frequency are the underlying cause of the previously observed synaptic changes. We observed that miniature synaptic recordings partially recapitulated the changes observed with spontaneous synaptic recordings. In slices from PE females, we found that aVMHvl^Pr+^ neurons received a significantly higher frequency of mIPSCs compared to the frequency of events sensed by PR+ neurons from D females (aVMHvl PE median [IQR]=2.06 [1.40 3.08] Hz; D=0.73 [0.65 0.99] Hz; 2-way ANOVA, reproductive phase x AP interaction factor, F(2,59)=6.24, p=0.003; Sidak’s multiple comparison test, anterior PE vs. D p=0.001). In contrast, the robust decrease of sEPSCs observed in the receptive phase for anterior and medial VMHvl^Pr+^ neurons was visibly reduced in the presence of TTX, despite giving a small but significant decrease in the frequency of mEPSCs received by VMHvl^Pr+^ neurons in the PE phase compared to the D phase (2-way ANOVA, reproductive phase, F(1,59)=4.27, p=0.043), this effect no longer yielded significant differences when correcting for multiple comparisons for anterior or medial neurons (Fig. S6E). While the amplitude of mEPSCs remained unchanged across the reproductive cycle, there was an observable increase in the amplitude of mIPSCs recorded from aVMHvl^Pr+^ neurons obtained from PE females, but also not large enough to yield statistical significance (Fig. S6H).

The changes in the frequency of miniature synaptic events yielded a significant shift in the E/I frequency balance of aVMHvl^Pr+^ neurons across the female reproductive cycle (aVMHvl PE median [IQR]=0.17 [–0.14 0.41] a.u.; D=0.69 [0.48 0.83] a.u.; 2-way ANOVA, reproductive phase x AP interaction factor, F(2,59)=3.97, p=0.024; Sidak’s multiple comparison test, anterior PE vs. D p=0.002), in line with the previous results (Fig. S6I), but in this case largely dominated by an increase in inhibitory transmission, which indicates that the modulation of presynaptic firing plays a sizeable role in the excitatory synaptic frequency changes observed across the female’s reproductive cycle.

Next, to determine if presynaptic release probability was responsible for the changes in the frequency of synaptic inputs, we measured the paired pulse ratio (PPR) of electrically evoked synaptic responses (see methods). To our surprise, we observed a small but significant PPR decrease of the paired evoked excitatory responses (eEPSCs) of VMHvl^Pr+^ neurons from PE females showed short term depression, which was not observed in PR+ neurons from D females (Fig. S6J; 2-way ANOVA, reproductive phase, F(1,80)=8.52, p=0.0045). The decrease in eEPSCs-PPR, which indicates an increase in presynaptic release probability, was apparent in anterior and medial neurons, however only the former showed significant differences after correcting for multiple comparisons (aVMHvl^Pr+^ PE median [IQR]=0.75 [0.62 0.98] a.u.; D= 0.94 [0.87 0.99] a.u.; 2-way ANOVA; Sidak’s multiple comparison test, anterior PE vs. D p=0.02). This seemingly paradoxical increase in release probability of PE aVMHvl^Pr+^ excitatory inputs maybe explained within the depletion model of synaptic transmission, by a largely intact readily releasable pool of synaptic vesicles given a much-reduced basal transmission during PE phase (Fig. 2B).

In contrast, the PPR of inhibitory responses (eIPSCs) did not undergo any significant modulation across the female reproductive cycle at any AP level of the VMHvl (Fig. S6K). However, it is worth noting that, when comparing the relative amplitude of the eEPSCs and eIPSCs, we observed a significant E/I balance shift in aVMHvl^Pr+^ neurons (Fig. S6L; aVMHvl PE median [IQR]=0.15 [–0.08 0.51] a.u.; D=0.46 [0.39 0.57] a.u.; 2-way ANOVA, reproductive phase x AP interaction factor, F(2,160)=4.96; p=0.0093; Sidak’s multiple comparison test, anterior PE vs. D p=0.0058), consistent with our observations when recording spontaneous synaptic events.

Finally, to dissect the origin of the changes in the inhibitory synaptic input sensed by aVMHvl^Pr+^ neurons, we made use of neurotransmitter uncaging^34^ . We recorded synaptic currents evoked by the light-dependent uncaging of GABA throughout the dendritic arborization of VMHvl^Pr+^ neurons from PE and D females using galvo mirrors to target a grid of laser stimulation locations (Fig. 3A–B, see methods). Replacing the presynaptic compartment, here mostly silent by the presence of TTX, by direct photo-release of GABA enabled us to directly test changes originating at the postsynaptic level. In these conditions, changes on the amplitude of inhibitory currents evoked by GABA uncaging, together with the previously observed changes in frequency but not amplitude of spontaneous synaptic events, is indicative of an increase in active synaptic sites, as a higher number of synapses may be activated by the same amount of photo-released GABA. We observed that the photo-release of GABA elicited significantly larger inhibitory currents in aVMHvl^Pr+^ neurons from PE females. This increase was restricted to the perisomatic dendritic domain of aVMHvl^Pr+^ neurons (Fig. 3C,E; <200 µm from the soma, aVMHvl^Pr+^ PE median [IQR]= 443.25 [349.26 583.72] pA; D=276.34 [151.79 332.69] pA; 2-way ANOVA reproductive phase factor, F(1,40)=4.87; p=0.03; Sidak’s multiple comparison test, anterior PE vs. D p=0.009) as it was absent both in the distal dendritic domains (>200 µm, Fig. 3D) as well as in medial and posterior VMHvl^Pr+^ neurons (Fig. 3C–G).

**Figure 3.**
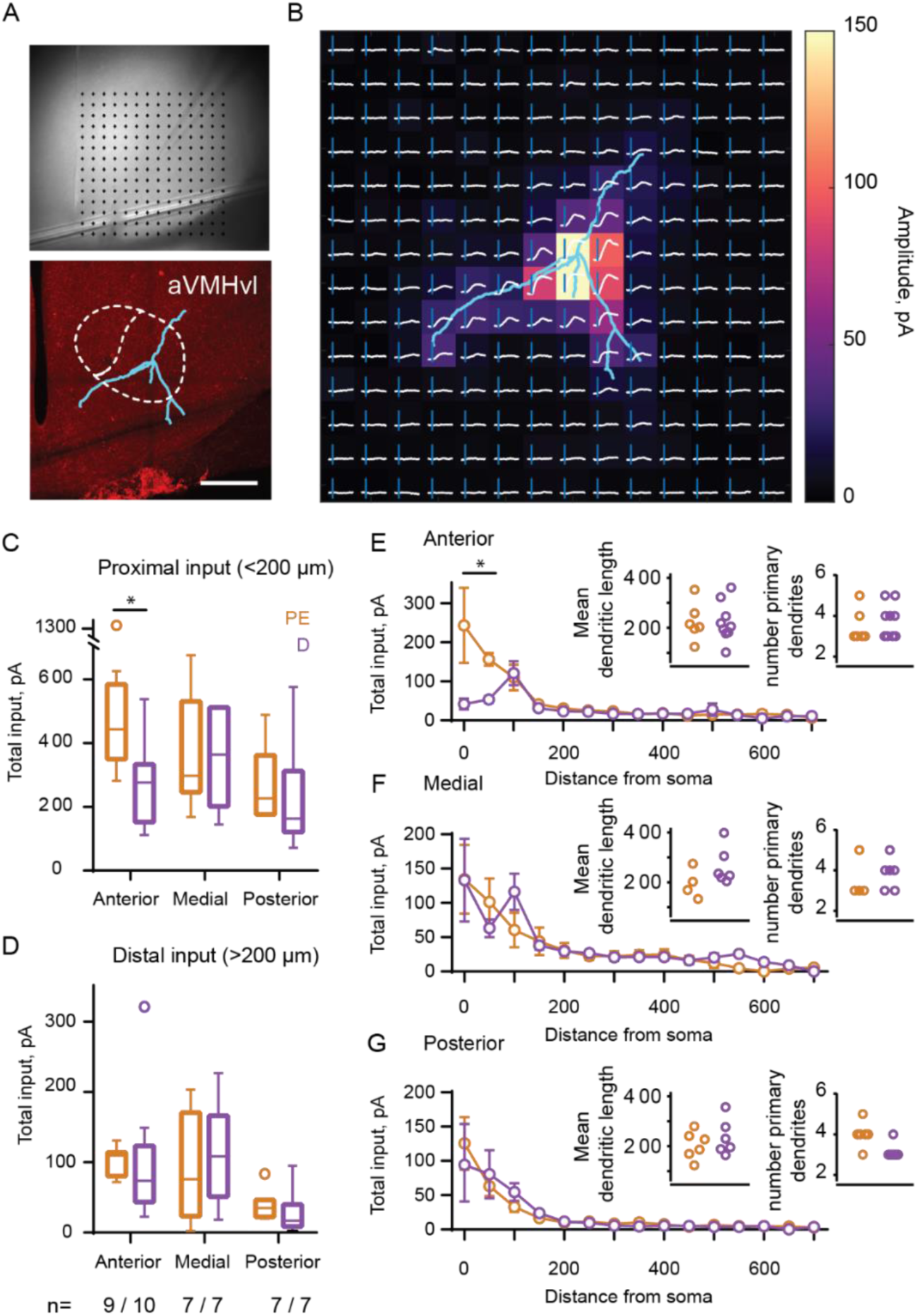
Uncaging of GABA reveals stronger perisomatic inhibitory responses in aVMHvl^Pr+^ neurons from PE females. (A) Representative example of a PR+ neuron recorded in the aVMHvl. Top picture represents the location of the recording pipette in the electrophysiology setup and the grid of laser stimulation locations. Bottom picture shows the post-fixation reconstruction of the neuron and its location within the aVMHvl. Scale bar: 400 µm. (B) Neuron reconstruction aligned to the uncaging GABA responses represented both as electrophysiological traces and as a heatmap of their peak amplitude (C-D) Total synaptic input (sum of active pixels, see methods) for the proximal (C) and distal (D) dendritic domains of anterior, medial and posterior VMHvl neurons from D and PE females. The n represents the number of neurons per recording condition. * represents p<0.05 using Sidak’s multiple comparisons test (E-G) Synaptic input amplitude as a function of dendritic distance from the soma and mean dendritic length and number of primary dendrites for anterior (E), medial (F) and posterior (G) VMHvl neurons from D and PE females. * represents p<0.05 using Sidak’s multiple comparisons test.

Importantly, these results suggest that the reproductive state correlated changes in inhibitory synaptic input may be due to cyclical modulation of the number of perisomatic inhibitory synapses onto aVMHvl^Pr+^ neurons.

Overall, our characterization of the synaptic input onto VMHvl^Pr+^ neurons across the female reproductive cycle draws a complex landscape with multiple forms of synaptic plasticity, mostly present in the anterior portion of the VMHvl, of presynaptic origin for excitatory inputs, and of postsynaptic origin for inhibitory ones. Importantly, these changes in synaptic input strongly correlate with the previously described activity changes of aVMHvl^Pr+^ neurons *in vivo*.

### Optogenetic activation of aVMHvl^Pr+^ neurons is sufficient to drive fighting behavior in PE females

Our results so far indicate a correlative relationship between the activity of aVMHvl^Pr+^ neurons and the expression of sexual fighting behavior in D females. To establish causality and assess whether activating this population could override the increase in sexual receptivity (or decrease in fighting) that occurs during the receptive phase of the reproductive cycle, we performed *in vivo* optogenetic activation of aVMHvl^Pr+^ neurons in PE females during a sexual behavior assay. For this, we bilaterally injected PR-Cre females with Cre-dependent AAV vectors expressing either ChR2-EYFP (ChR2 group) or EYFP (control [Ctrl] group) in the anterior portion of the VMHvl and implanted fiber cannulas for light delivery (Fig. 4A). After at least 2 weeks of recovery, we monitored the behavior of the ChR2 and Ctrl groups in open field and urine preference tests followed by a sexual behavior assay (Fig. 4B). Both groups of females, ChR2 and Ctrl, were PE, determined by a vaginal smear (performed before and in some cases also after the behavioral testing, see methods for details). Importantly, our stimulation pattern elicited a robust increase in the expression of cFos in the aVMHvl^Pr+^ population of ChR2 but not Ctrl females *in vivo* (Fig. 4C,D; Ctrl median [IQR]=7.49 [2.7 13.2] AAV+cFos+/AAV+ vs ChR2 median [IQR]=54.93 [37.5 70.8] AAV+cFos+/AAV+; Mann–Whitney U test, p=0.003). Consistently, aVMHvl^Pr+^ neurons were able to track the 20Hz optogenetic stimulation with high fidelity *ex vivo* (Fig. 4E).

**Figure 4.**
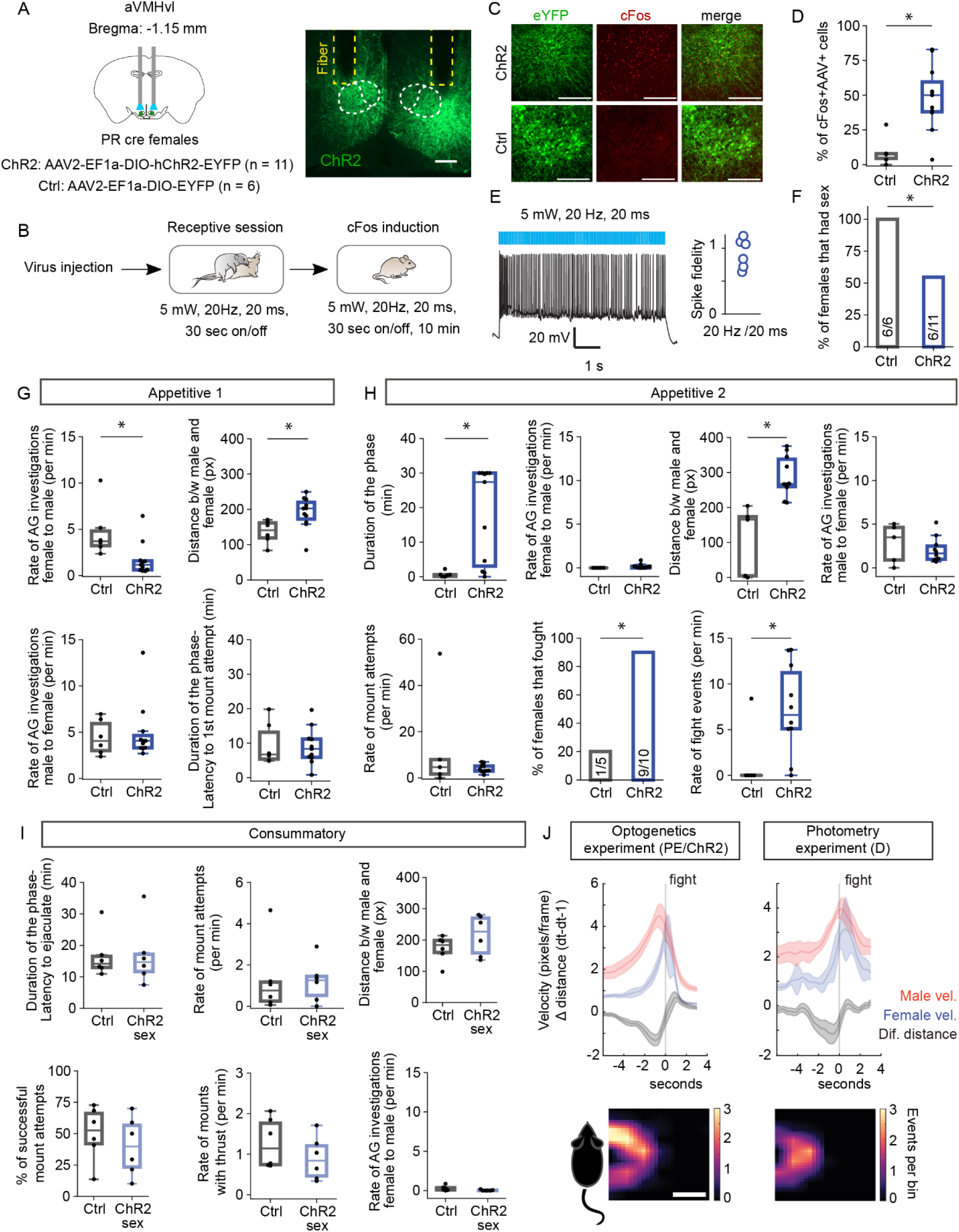
Optogenetic activation of aVMHvl^PR+^ neurons lead to an increase in fighting behavior. (A) Viral constructs and implantation coordinates. Representative image showing ChR2 expression and fiber placement. Scale bar: 200 μm. (B) Schematic illustrating the experimental design. (C) Representative images showing ChR2, cFos and double positive cells (top) or EYFP, cFos and double positive cells in ChR2 and Ctrl females, respectively. Scale bars: 100 μm. (D) Quantification of percentage of AAV+ cells that are double positive for cFos in Ctrl versus ChR2 females. * represents p<0.05 using Mann-Whitney U test. (E) Left: Representative trace from a ChR2 transfected aVMHvl^Pr+^ neuron recorded in current clamp mode and being stimulated with 20 ms pulses of blue light (5mW) at 20Hz for 5 seconds. Right: Spike fidelity (spikes measured/ number of light pulses delivered) for the individual aVMHvl^Pr+^ neurons recorded. (F) Percentage of females that had sex within 30 minutes after the first mount attempt by the male.* represents p<0.05 using the Z test. (G). Comparison of various behaviors displayed by Ctrl and ChR2 females during the appetitive 1 phase. * represents p<0.05 using Mann-Whitney U test. (H). Comparison of various behaviors displayed by Ctrl and ChR2 females during the appetitive 2 phase. 1 Ctrl and 1 ChR2 female did not have an App2 phase, i.e., the first mount attempt led to a successful mount with thrusts. * represents p<0.05 using Mann-Whitney U test. (I). Comparison of various behaviors displayed by Ctrl and ChR2 females during the consummatory phase. * represents p<0.05 using Mann-Whitney U test. (J) Top: Velocity profiles for the male (red) and female (blue) and the corresponding difference in distance over time between the couple (black) around a fight event (solid line) for females in the ChR2 group and D females from the photometry experiment. Shaded error bars indicate mean and standard error across mice. Bottom: Binned distribution of male locations with respect to the female at the moment of fight.

In line with our prediction, a significantly lower fraction of aVMHvl^Pr+^ ChR2 females engaged in sexual behavior when compared to Ctrl females (6 out of 11 vs 6 out of 6; Fig. 4F; Z test, p=0.027), suggesting that aVMHvl^Pr+^ activation may lead to an increase in fighting behavior displayed by PE females and thus fewer individuals allow the male to copulate with them.

To further characterize the effect of optogenetically stimulating the aVMHvl^Pr+^ population on the social interaction, we compared the behavior of Ctrl and ChR2 females during the three phases of sexual behavior — App1, App2 and Cons (Fig. 1A). In App1, ChR2 females displayed a lower rate of male-directed anogenital investigations when compared to Ctrl females (Fig. 4G; Ctrl median [IQR]=3.71 [2.93 5.44] events per minute vs ChR2 median [IQR]=1.17 [0.61 1.64] events per minute; Mann–Whitney U test, p=0.01) and there was on average higher distance between the couple (Ctrl median [IQR]=140.8 [108.5 165.5] pixels vs ChR2 median [IQR]=202.6 [163.2 228.3] pixels; Mann–Whitney U test, p=0.015). The lower rate of male-directed investigation and higher distance between the couple observed in the ChR2 group does not seem to derive from a lack of interest from the male, evidenced by comparable rates of anogenital investigation towards the female and latency to perform the first mount attempt in both groups (Fig. 4G).

Considering a large percentage of ChR2 females never accepted copulation attempts and remained in App2, the duration of this phase was naturally longer in the ChR2 group than in the Ctrl group (Fig. 4H; Ctrl median [IQR]=0.50 [0.03 1.01] minutes vs ChR2 median [IQR]=27.39 [1.52 30] minutes; Mann–Whitney U test, p=0.007). Interestingly, the transition to App2 is characterized by a drastic decrease in the rate of anogenital investigation that Ctrl females display towards the male (Fig. 4G-H). This does not only support the division of the App phase into two sub-phases (App1 and App2), but also may explain the similar rates of anogenital investigations of the two groups after the male-initiated copulation attempts due to a floor effect (Fig. 4H). The distance between the couple in the ChR2 group remained higher during App2 (Fig. 4H; Ctrl median [IQR]=165.2 [2.31 189.1] pixels vs ChR2 median [IQR]=267.4 [247.5 349.8] pixels; Mann– Whitney U test, p=0.0007). During App2 males from the two groups investigated and attempted copulation at similar rates (Fig. 4H), further supporting the conclusion that the disruption in the social interaction is caused by the female and not by a decreased motivation of the males interacting with the ChR2 females.

App2 is characterized by male mount attempts, which may lead to strong rejection behavior in the form of fights (boxing and kicking) by D females (Fig. 1A). Upon examining female rejection behavior during this phase we found that a greater percentage of ChR2 females displayed fighting behavior compared with Ctrl females (9 out of 10 vs. 1 out of 5; Z test, p=0.003), at a very high rate (Fig. 4H; Ctrl median [IQR]=0 [0 0.1] events per minute vs ChR2 median [IQR]=6.60 [3.96 12.44] events per minute; Mann–Whitney U test, p=0.029). These results show that the activation of the aVMHvl^Pr+^ population leads to elevated fighting behavior. Interestingly, the difference in fighting behavior observed during App2 when comparing ChR2 to Ctrl females is similar to the one observed when comparing the rejection behavior of D and PE females from our photometry experiments (Fig. S3B), suggesting that the artificial activation of aVMHvl^Pr+^ neurons shifts the behavior of PE females towards the behavior displayed by D females.

ChR2 females had a significantly longer App2 phase compared to Ctrl females, to ensure that the fighting behavior shown by the ChR2 group was consistently displayed as soon as the male tried to copulate instead of emerging after a prolonged period of mount attempts, for which a longer App2 phase may lead to an artifactual increase indirectly dependent on the optogenetic stimulation, we performed a random subsampling of App2 data from all animals that had a longer App2 than a set threshold based on Ctrl App2 durations (see methods). We found that behavioral differences are maintained in this random subsampling analysis (Fig. S7A–C). For both App1 and App2 we also compared several behavioral parameters between “light on” and “light off” periods and found no significant differences, suggesting that light stimulation did not acutely affect behavior (Fig. S7D–G). The behavioral differences between Ctrl and ChR2 females are also not caused by abnormalities in locomotion, anxiety or social cue processing in manipulated females as assessed in the open field test and a male/female urine preference test (Fig. S8). Finally, for the ChR2-No sex females we performed an additional vaginal smear at the end of the behavioral assay to ensure that the outcome of the experiment was not due to an error in the assessment of the reproductive stage nor because females had exited the PE phase. The vaginal smear of ChR2-No sex females looked like the smear of PE females (data not shown).

We next compared the behavior of the subset of ChR2 females that had sex (ChR2 sex) to Ctrl females during the Cons phase. Interestingly, we did not find notable differences between the two groups in any of the parameters that differed between manipulated and Ctrl females during App1 and App2, such as the distance between the male and female, the rate of investigations between them and rate of mount attempts displayed by the male (Fig. 4I). The percentage of mount attempts that resulted in successful mounts (Fig. 4I) was also comparable, pointing to the possibility that once copulation is initiated, the fighting behavior resulting from the artificial stimulation of aVMHvl^Pr+^ neurons is not sufficient to disrupt the sexual interaction anymore.

In summary, the optogenetic activation of the aVMHvl^Pr+^ population caused a significant increase in the display of fighting behavior by PE females, which in some cases was sufficient to block copulation. For the ChR2 females that allowed copulation, once the Cons phase was initiated, no obvious difference to Ctrl females was observed.

Given the well-established involvement of the VMHvl in maternal aggression^25^, we next sought to further inspect the fighting behavior displayed by ChR2 females, in particular if fighting behavior was initiated by the female or a reaction to male approach or mount attempts, and whether the nature of this fighting behavior varies across different treatment groups and experiments. For this, we compared several aspects of the female’s (and the couple’s) behavior for the ChR2 and Ctrl groups, alongside that of D and PE couples from the photometry experiments using neural networks–assisted pose estimation algorithms^35^.

To assess if fighting was initiated by the female, we inspected the behavior of the couple in the seconds preceding their execution. For that we aligned the first boxing or kicking event from a bout of fighting events and determined the velocity of each animal and the distance between the couple. Our analysis revealed that in all cases, just before a fight event, the male’s velocity began increasing first, concomitant with a reduction in the distance between the two animals, indicating that the male was the one moving towards the female (Fig. 4J). This was closely followed by an increase in the velocity of the female coupled with the distance between the animals increasing once again, indicative of the movement of the female away from the male. This analysis rules out the question of aggression, showing that boxing and kicking events are not sought by females but rather occur in response to male pursuit. The location where fight events took place relative to the position of the female also remains similar across groups (Fig. 4J). All other behavioral metrics, except for the center occupancy of the behavioral box during App1 (Fig. S9) remained unaltered during the social interaction, supporting the specificity of the neuronal manipulation.

Overall, these results uncover a hitherto unknown role of aVMHvl^Pr+^ in driving sexual fighting behavior, corroborating our hypothesis that a higher activity of these cells during the non-receptive phase may be responsible for high rejection rates, while a suppression in their activity during the receptive phase allows for low rejection rates and successful engagement in sexual acceptance behavior.

## Discussion

In the present study, we have identified a novel role for the aVMHvl in the cyclical control of rejection behavior. We find that PR+ neurons in the aVMHvl are more active during the non-receptive phase of the reproductive cycle, in which the female vigorously defends herself from unwanted copulation attempts, and conversely, their lower activity correlates with a reduced rejection rate that is permissive to sexual behavior to occur during the receptive phase. These internal state–dependent differences in aVMHvl activation likely reflect the higher engagement of aVMHvl^Pr+^ neurons, which are highly tuned to fighting behavior during the non-receptive phase. Furthermore, aVMHvl^Pr+^ neurons undergo a shift in E/I synaptic balance throughout the reproductive cycle, being lower during the female’s receptive period and originating in both pre- and postsynaptic compartments. Crucially, optogenetic activation of aVMHvl^Pr+^ neurons during the receptive phase of the reproductive cycle was able to disrupt behavioral receptivity by increasing fighting rejection behavior. These findings not only suggest that the cyclical control of fighting behavior is exerted by synaptic changes that silence “fighting” tuned aVMHvl^Pr+^ neurons in the sexually receptive phase, but also expand our knowledge on the spatial distribution of functions carried out by the VMHvl in the female brain.

Previous studies have shown that mediolaterally segregated ensembles of neurons in the pVMHvl were responsible for mating and for female aggression^12^, the occurrence of the latter being restricted to specific moments of the female’s life, such as during the lactating phase of motherhood^25^. Here, we provide evidence that instead of promoting active aggression, the activation of aVMHvl^Pr+^ neurons promotes self-defense in response to undesired copulation attempts. It is worth noting that, in male mice, the aVMHvl is active upon conspecific aggression self-defense^24^. Thus, the emerging picture points to the aVMHvl as a putative hub for the control of conspecific self-defense against undesired actions of diverse nature in both sexes.

In contrast, increasing evidence has consolidated the role of the pVMHvl in the control of female mating, showing that several forms of neural plasticity are crucial for its control and expression throughout the reproductive cycle^2, 5, 13, 25^. Previous studies have shown that the output terminals of pVMHvlPr+ neurons to the AVPV undergo structural potentiation in response to increasing levels of sex hormones in a process that is likely crucial for its facilitatory role on mating^2^. Furthermore, it has been recently shown that a subpopulation pVMHvl neurons expressing the cholecystokinin receptor (Cckar) undergoes synaptic changes supporting an increased mating drive in receptive females^13^. These changes entail an increased E/I synaptic balance during the female receptive phase, which confirms previous reports using immunochemical detection of markers of synaptic potentiation such as the GluA1 AMPA receptor^20^. In our data, we were unable to identify synaptic potentiation in pVMHvl^Pr+^ neurons; however, as reported by Yin and colleagues, these changes were restricted to a relatively small subpopulation of pVMHvl neurons (Cckar+) and, thus, likely masked in our recording conditions using a broader genetic handle such as the PR.

Importantly, the mating-promoting synaptic changes described for pVMHvl^Cckar+^ neurons are in the opposite direction of what we report for aVMHvl^Pr+^ fighting cells. Altogether, these findings suggest a working model in which sex hormones simultaneously orchestrate a reduction of defensive/rejection tendencies by lowering E/I in the aVMHvl, while increasing the mating drive by increasing E/I in the pVMHvl of females during the fertile/sexually receptive phase to allow sexual behavior to occur. Furthermore, our optogenetics results show that the activation of aVMHvl^Pr+^ neurons was sufficient to significantly decrease the percentage of females in the reproductive phase of the cycle engaging in sexual behavior, which suggests that this decrease in self-defensive tendencies is a crucial permissive step for the natural expression of female sexual behavior.

In the present study, we provide for the first time, a characterization of the activation profile of aVMHvl^Pr+^ neurons performing a socio-sexual assay and compare it to that of pVMHvl^Pr+^ neurons in female mice throughout the reproductive cycle of naturally cycling females. We observed convergent and divergent motifs of activation of these subareas during different behaviors. Neurons in the pVMHvl^Pr+^ were responsive to male chemosignals and copulation. This is consistent with previous research reporting pVMHvl activity during male investigations and successful mounts, including the moment in which the male initiates the mount (stimulating the back flanks of the female) but also during the display of the lordosis posture for maintaining copulation^2, 13^. However, whereas aVMHvl^Pr+^ neurons were also active upon male conspecific sensing, they were inhibited during the initiation and maintenance of the mount. And as opposed to their posterior counterparts, aVMHvl^Pr+^ neurons were active during the display of fighting behavior by non-receptive females.

Recent reports have highlighted the heterogeneity of the VMHvl throughout its AP axis from the transcriptomic, anatomic and physiological perspectives^18, 22, 23^. Whereas the pVMHvl preferentially produces afferents to hypothalamic nuclei, the aVMHvl seems to preferentially project to the midbrain periaqueductal gray^22, 24^. Furthermore, although the projection pattern of VMHvl neurons does not appear to present sexual dimorphism^22^, the density of PR+ neurons in the VMHvl presents dimorphic differences, being higher in females than males^11^. Future studies are necessary to shed light on the downstream projections of the aVMHvl^Pr+^ population and determine their specific contribution to the expression of fighting behavior and to understand whether aVMHvl^Pr+^ neurons participate as a whole in the control of rejection, or if there are genetically defined functional subclusters as it has been shown for the pVMHvl^12, 13^.

Here, we provide evidence supporting the coexistence of a presynaptic decrease in basal excitatory transmission with mechanisms of postsynaptic potentiation of inhibitory synapses that possibly act in synergy to decrease the activity of aVMHvl^Pr+^ during the sexually receptive phase. While the role of distributed synergistic plasticity is well established in other neural systems^36^, in the hypothalamus we are only beginning to understand the mechanistic origin and function of synaptic plasticity processes^5, 13, 37, 38^. In males, a form of experience- and testosterone-dependent LTP mediates aggression learning^38^ and likely participates in the changes in neural representations occurring in the male brain after aggression^39^. However, in the female brain, the emerging picture seems to indicate that synaptic plasticity can support cyclic changes in the female sexual brain even in the absence of any sexual experience (i.e. naive females). This form of cyclic plasticity suggests that sex hormones may orchestrate multiple forms of plasticity throughout the female brain for the neural control of sexual behavior^2, 5, 13^. From a mechanistic perspective, several candidate effector pathways could modulate synaptic transmission in response to fluctuating levels of sex hormones. Retrograde messengers, such as nitric oxide (NO), can diffuse from the post- to the pre-synaptic compartment where they can exert modulatory actions on synaptic transmission^40^. NO-synthesizing neurons in the VMHvl are fundamental for the expression of sexual receptivity^9^, consistent with the fact that pharmacological blockade of NO synthesis impairs female sexual behavior^41^. However, while a cluster of NO-synthesizing neurons is visible at the pVMHvl^42, 43^, it is unclear whether the aVMHvl is equipped with a similar neuronal ensemble, and thus, its relation with the plastic processes here reported remains unclear. Furthermore, the VMHvl and neighboring hypothalamic nuclei possess an extensive repertoire of retrograde messengers, including opioids, cannabinoids and neurotrophic factors^44–46^, most of which have been shown to be involved in other hypothalamic dependent behaviors, such as feeding, and whose involvement in sexual behavior relevant synaptic plasticity is yet to be explored and should be focus of future research. Finally, a considerable number of areas projecting to the VMH are themselves equipped with sex hormone receptors that could, in parallel, give origin to the observed changes in presynaptic firing as it has been previously shown for areas the medial preoptic hypothalamus to medial amygdala pathway^55^. However, it remains to be determined whether these changes in basal transmission play a significant role modulating the activity of aVMHvl^PR+^ neurons in vivo.

Another major form of neural plasticity recently described in the context of sexual behavior, is the structural potentiation of axonal pathways. This has been recently shown to occur in the pVMHvl→AVPV afferents in response to increasing levels of sex hormones and produced, as a readout, an increase in the frequency of synaptic currents in the postsynaptic AVPV neurons^2^. Interestingly, we observe a similar increase in IPSC frequency recorded from aVMHvlPr+ neurons in the receptive phase of the female. These observations suggest that a possible mechanism underlying the form of inhibitory plasticity here reported is the structural potentiation of one or more of its inhibitory incoming pathways, a hypothesis that should be tested in future work.

Despite the fact that some mechanistic aspects of the synaptic plasticity processes observed remain unanswered, our findings contribute to reconciling previously seemingly contradictory bodies of literature. While it is becoming clear that the activity potentiation of neural subpopulations within the pVMHvl is necessary for the display of female sexual receptivity^2, 13^, classical pharmacological studies have shown that diminishing glutamatergic activity or increasing GABAergic tone in the VMHvl results in a paradoxical increase in female sexual receptivity^10, 47^. These pharmacological manipulations will result in a reduced E/I ratio that enhances female receptivity. In light of our findings, a lower E/I ratio in specific subpopulations of neurons mostly present in the anterior part of the nucleus, strongly correlates with female sexual receptivity and its exogenous activation is sufficient to significantly disrupt sexual behavior. Altogether, sexual receptivity seems to arise from the downregulation of rejection behavior and the upregulation of sexual receptivity through the bidirectional modulation of the E/I balance of anterior (rejection) and posterior (mating) neurons of the VMHvl. How the same signal (E and P) leads to opposing outcomes in these two populations of neurons begs further investigation.

In summary, with the present study we propose to update the current model of VMHvl function in the control of female sexual behavior. When females are non-receptive, sex is efficiently prevented via the combined action of the inability of the pVMHvl to facilitate lordosis plus the execution of rejection behavior driven by the aVMHvl^Pr+^ population. In contrast, when females are receptive, the decrease in E/I synaptic balance impinging on aVMHvl^Pr+^ neurons leads to the suppression of rejection behavior, allowing lordosis to be expressed via the potentiated input/output of pVMHvl neurons^2, 13^. Altogether, our results support a hierarchical organization of sexual behavior, where the downregulation of self-defense mechanisms is paramount for sexual receptivity to emerge during the receptive phase of the cycle. While our results strongly suggest that rejection behavior can indeed override sexual receptivity, as the optogenetic activation of aVMHvl^Pr+^ neurons increased rejection behavior (preventing copulation in half of the couples despite being in the most receptive phase of the cycle), mounting also led to a decrease in the activity of the aVMHvl^Pr+^ population, as see during the photometry recordings. In addition, once successful mounting is initiated, the artificial activation of the aVMHvl^Pr+^ population no longer affects copulation. This idea brings to mind the hierarchical organization of decisions proposed by Tinbergen^48, 49^, with rejection and mating consummatory acts viewed as parallel modules implemented by separate circuits which are controlled by a common reproductive hierarchy. In this model, engagement in each of the actions, rejection for example, lowers the probability of the individual executing the opposing action, mounting in this case, increasing the robustness of such decision strategy. Our study provides evidence that mounting behavior may also contribute to that robustness, indirectly inhibiting the aVMHvl^Pr+^ population (photometry), lowering the probability of rejection once copulation was initiated, and probably explaining why once sex starts, the anterior VMHvl does not affect copulation anymore (optogenetics).

## Acknowledgments

We thank the Lima lab for their critical input during the course of this project. This work was supported by the Champalimaud Foundation through Fundação para a Ciência e a Tecnologia (FCT) Portuguese national funds in the context of the project UIDB/04443/2020, by the research infrastructure CONGENTO, co-financed by Lisboa Regional Operational Programme (Lisboa2020), under the PORTUGAL 2020 Partnership Agreement, through the European Regional Development Fund (ERDF), and FCT under Project LISBOA-01-0145-FEDER-022170 (S.Q.L.), PD/BD/114305/2016 (B.F.A.H.), 2021.07274.BD (I.C.D.), and the European Research Council Consolidator Grant 772827 (to S.Q.L.).

## Author contributions

NGC, BFAH and SQL conceptualized this work and wrote the manuscript. NGC, BFAH, ICD, KN, MAD and LF conducted the research, collecting the data in here presented. NGC and BFAH analyzed the data and generated the visualization files presented in this study. BL adapted and developed software necessary for the data analysis included in this study. Funds were obtained by SQL.

## Supplementary figures

**Figure S1.**
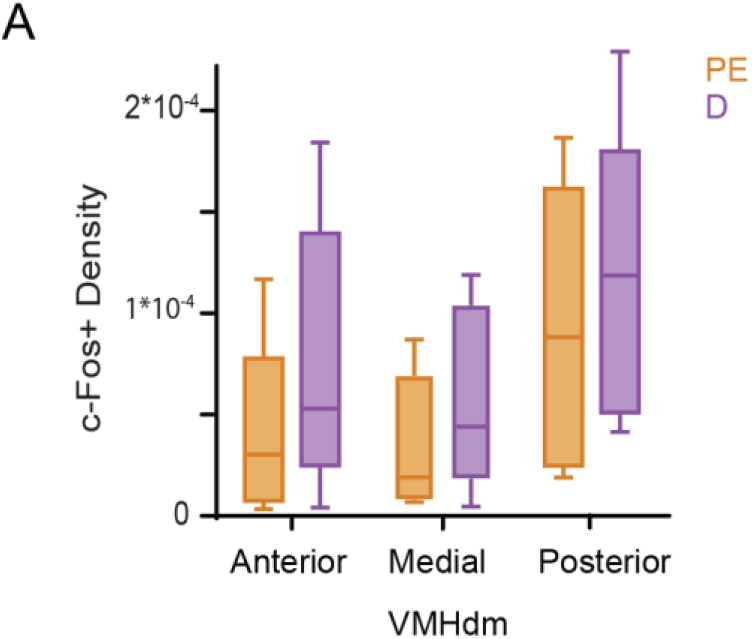
A similar pattern of activation is observed in the VMHdm of sexually rejecting and sexually accepting females. cFos-positive cell density in the VMHdm along the anterior-posterior axis in either proestrus/estrus (PE, orange) or diestrus (D, purple) females after sexual acceptance or rejection behavior, respectively.

**Figure S2.**
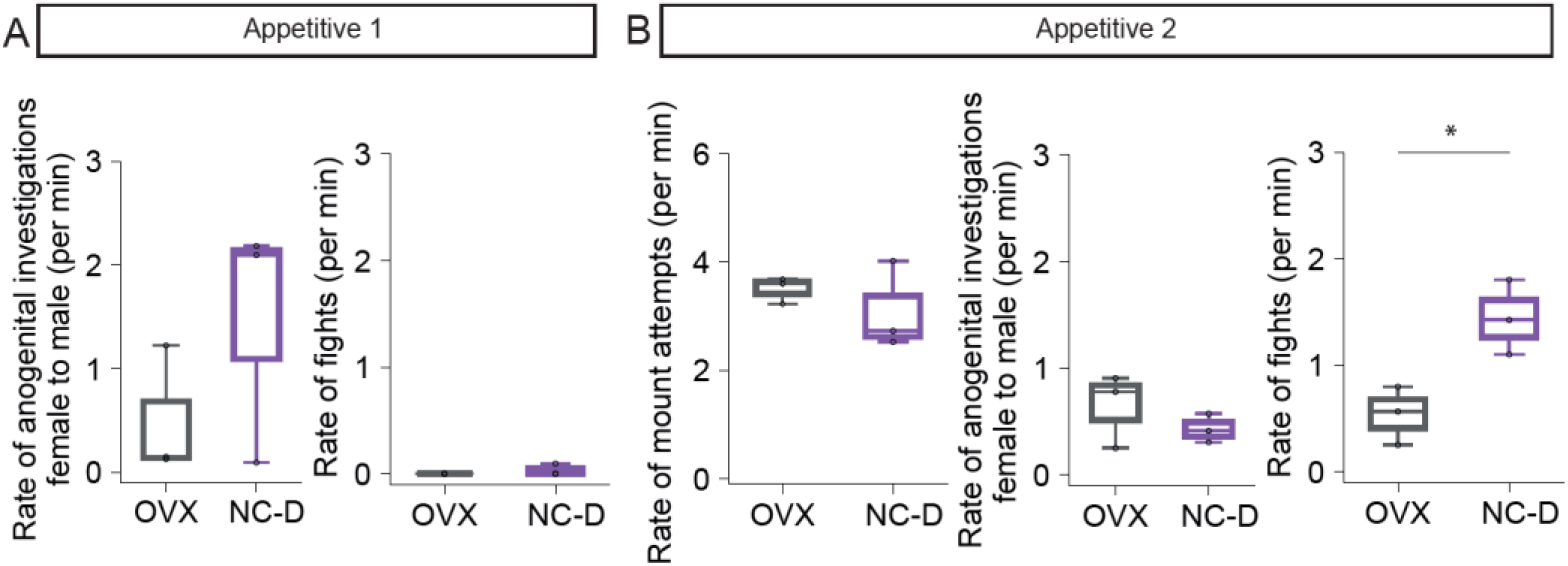
Non-primed ovariectomized females (OVX) display significantly lower amounts of fighting rejections compared to diestrus naturally cycling females (NC-D) in the appetitive phase 2. (A) Rate of male-directed anogenital investigations and fight rejections displayed by the female in App1. (B) Rate of mount attempts displayed by the male and rate of male-directed anogenital investigations performed and fight rejections displayed by the female in App2. * represents p<0.05 using unpaired t-test.

**Figure S3.**
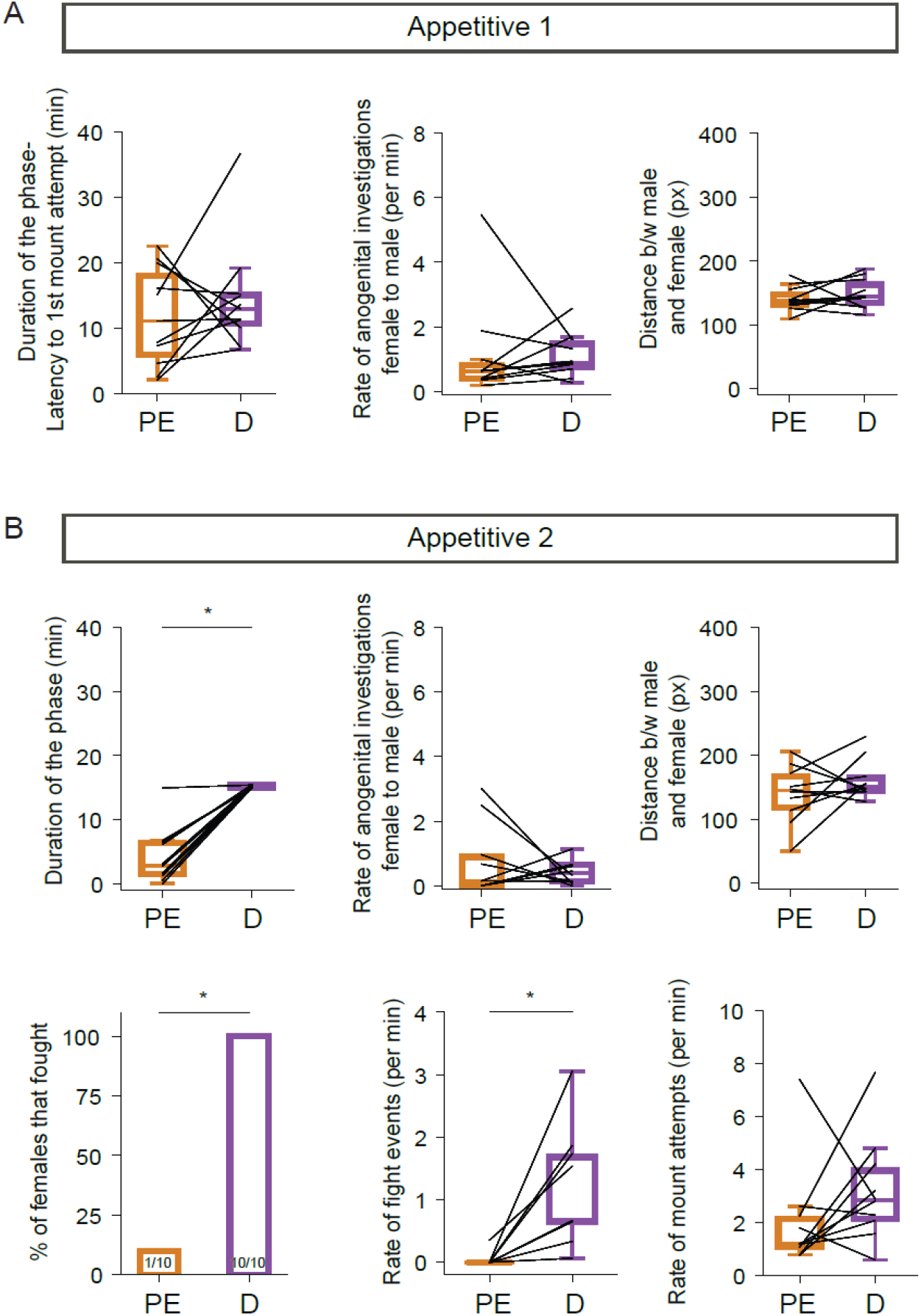
Behavior during the appetitive phases 1 and 2 of naturally cycling females during fiber photometry recordings. (A) Left: Duration of App1 which is the latency to the 1^st^ mount attempt from male entry into the arena. Center: Rate of male-directed anogenital investigations performed by the female in App1. Right: Average distance between the male and female during App1. (B) Top left: Duration of App2, which is the latency to the 1^st^ successful mount. * represents p<0.05 using Mann-Whitney U test. Top center: Rate of male-directed anogenital investigations performed by the female in App2. Top right: Average distance between the male and female during App2. Bottom left: Percentage of females that displayed at least 1 fight event in App2. * represents p<0.05 using the Z test. Bottom center: Rate of fight events displayed by females in App2.* represents p<0.05 using Mann–Whitney U test. Bottom right: Rate of mount attempts in App2.

**Figure S4.**
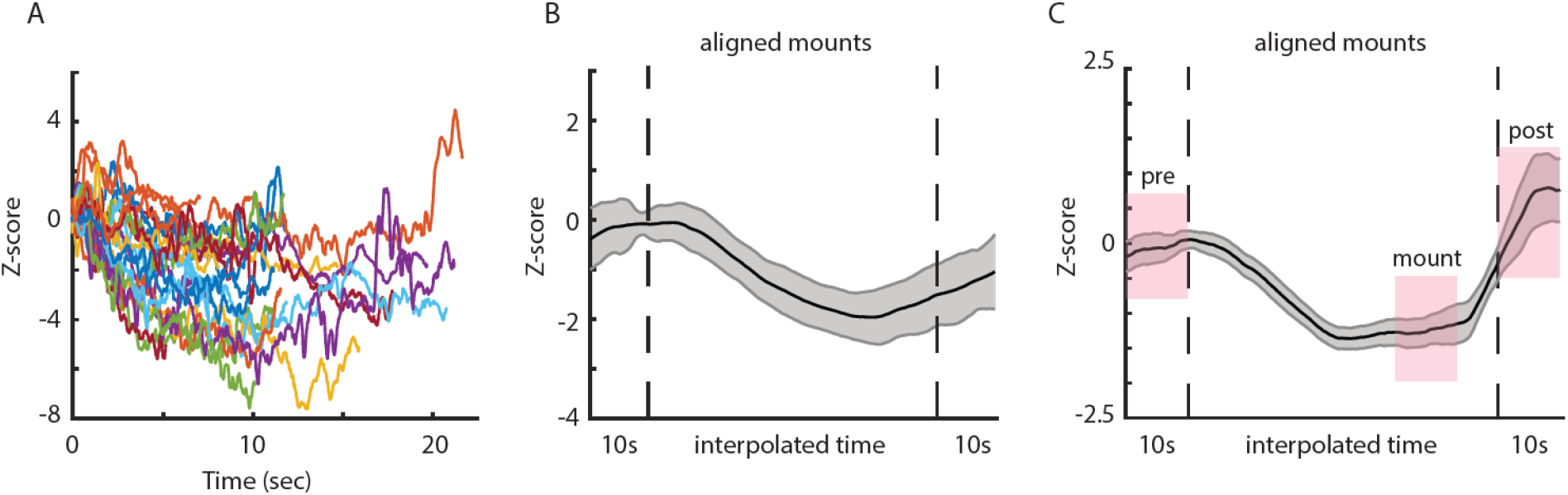
aVMHvl^Pr+^ neurons are inhibited throughout the full duration of the mount. (A) Photometry GCaMP signal from mount in to mount out of every individual mount of an example session. (B) Mounts in A were linearly interpolated to match the same time duration. Note the persistent reduction throughout the whole duration of the mounts. (C) Average of all linearly interpolated mounts for all the PE females of the photometry experiment. When comparing the pink shaded areas using a Mann–Whitney U test, we obtained: p= 0.0079 for the GCaMP response pre-mount vs mount, and p=0.15 for pre-mount vs post mount.

**Figure S5.**
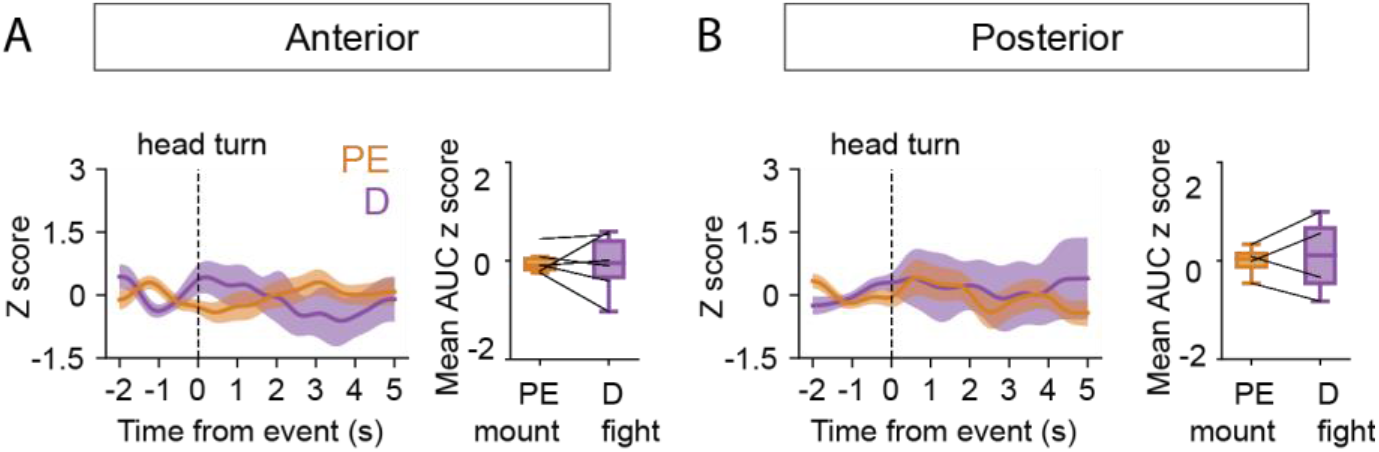
Head turn is not associated with a notable increase in fluorescence signal using fiber photometry. (A,B) PETHs showing z-scored head turn–associated GCaMP signal in anterior (A) and posterior (B) VMHvl^Pr+^ neurons. Boxplots comparing the area under the curve of the z score within 5 seconds after the event between non-receptive and receptive sessions. D: purple, PE: orange; p values obtained with a paired t-test.

**Figure S6.**
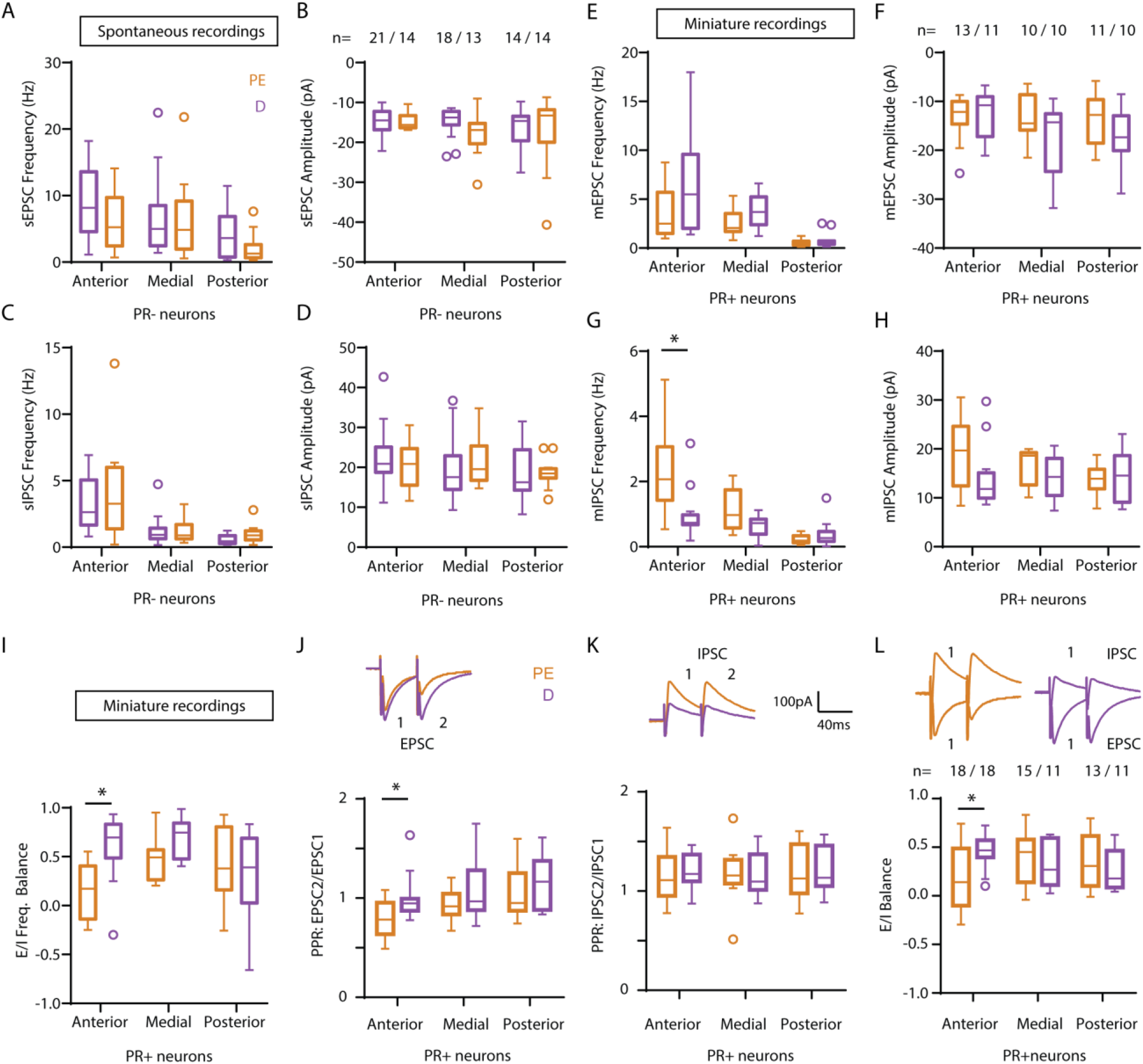
Spontaneous synaptic recordings of VMHvl.PR- and miniature recordings of VMHvl^Pr+^ neurons across the reproductive cycle. (A-D) Frequency and amplitude of excitatory and inhibitory spontaneous synaptic of VMHvl.PR-across the AP axis of the VMHvl and the reproductive cycle. (E-F) Frequency and amplitude of miniature excitatory synaptic events throughout the antero-posterior extension of the VMHvl in D and PE females in the presence of TTX. (G-H) Frequency and amplitude of miniature inhibitory synaptic events throughout the antero-posterior extension of the VMHvl in D and PE females in the presence of TTX. (I) Excitatory to inhibitory balance obtained from the frequency of miniature synaptic events. The n represents the number of neurons per recording condition, * represents p<0.05 using Sidak’s multiple comparisons test. (J) Paired pulse ratio of electrically evoked excitatory currents underwent a significant decrease in aVMHvl^Pr+^ neurons in PE compared to D. (K) Paired pulse ratio of electrically evoked inhibitory currents remained unchanged. (L) Furthermore, the relative change in electrically evoked responses yielded a significant E/I balance decrease in aVMHvl^Pr+^ neurons in PE compared to D. The n represents the number of neurons per recording condition, * represents p<0.05 using Sidak’s multiple comparisons test.

**Figure S7.**
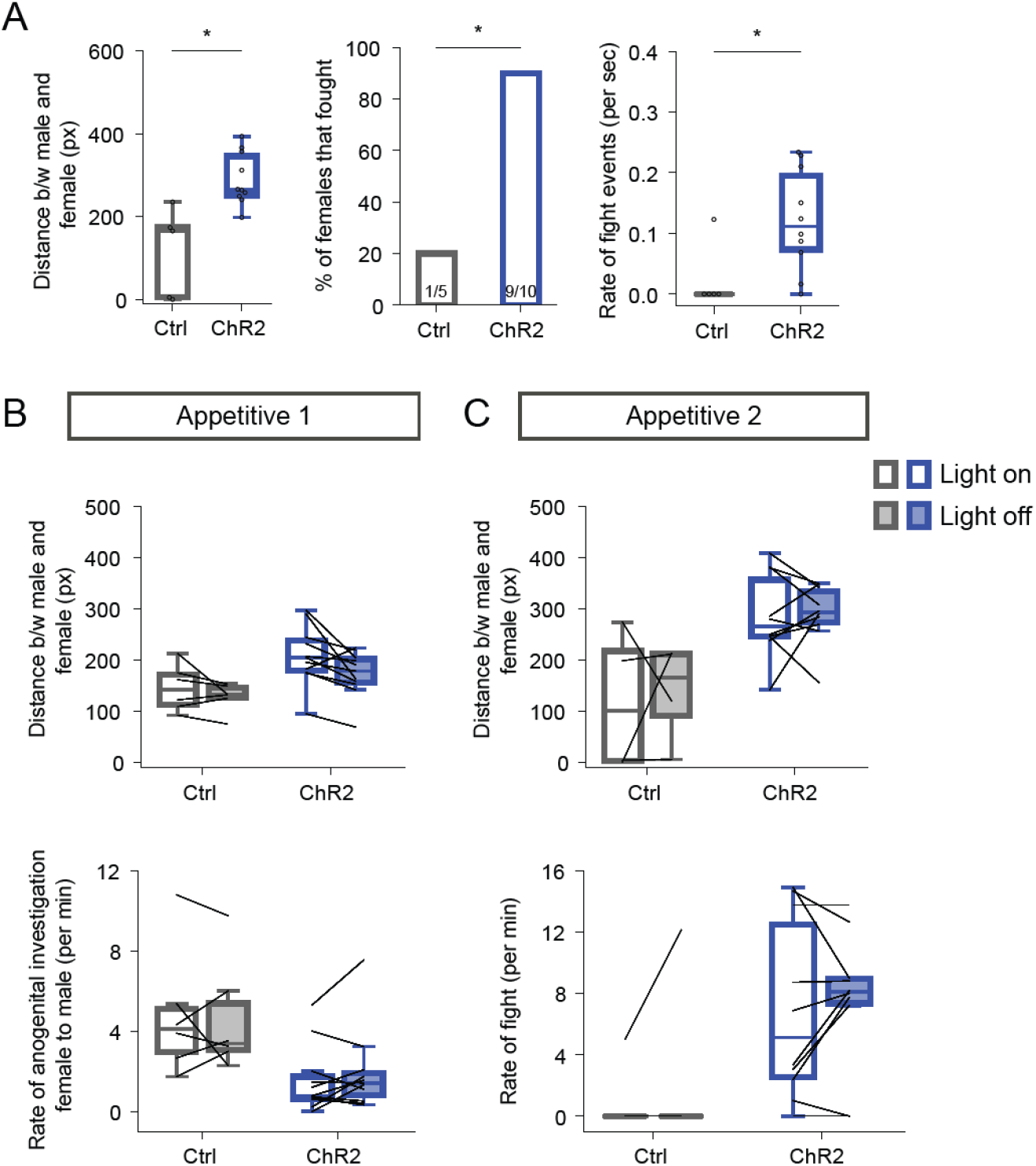
Effects of aVMHvl^Pr+^ stimulation on behavior are not affected by duration of App2 and are not acutely affected by light. (A) Distance between the male and female, percentage of females that displayed at least one fight event during the phase and rate of fight events displayed by females when a random 30-second window is used for all animals that had App2 duration >60 seconds. _(B-C)_ Average distance between the male and female and rate of male-directed anogenital investigations performed by the female during the ‘light on’ and ‘light off’ periods throughout appetitive 1 (B) and 2 (C) phases. B. Average distance between the male and female during the ‘light on’ and ‘light off’ periods of the phase. * represents p<0.05 using Mann– Whitney U test.

**Figure S8.**
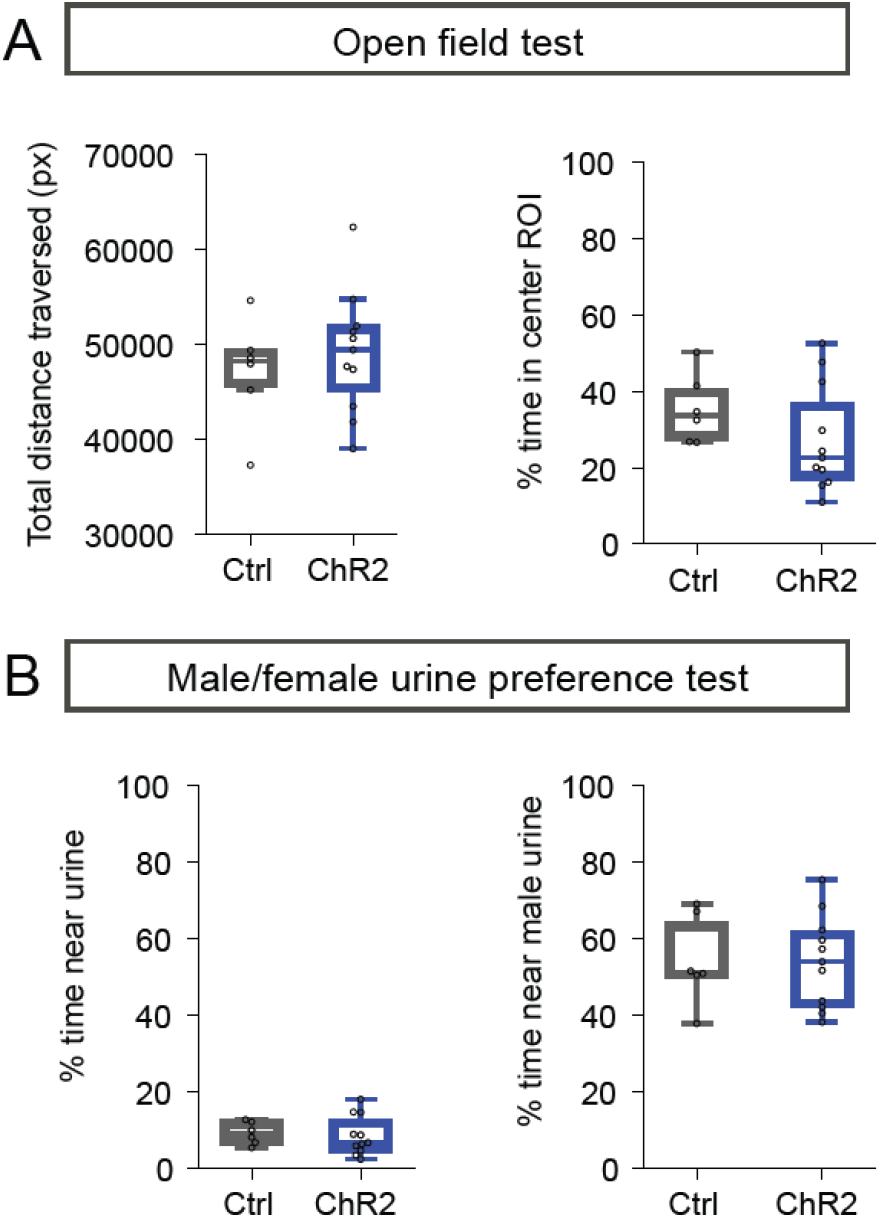
Optogenetic stimulation of aVMHvl^Pr+^ neurons does not lead to abnormalities in locomotion, anxiety or social cue interaction. (A) Left: total distance traversed by females in the open field; right: percentage of time spent by females in a central zone of the open field. (B) Left: percentage of time spent close to urine patches; right: percentage of time spent close to the male urine patch.

**Figure S9.**
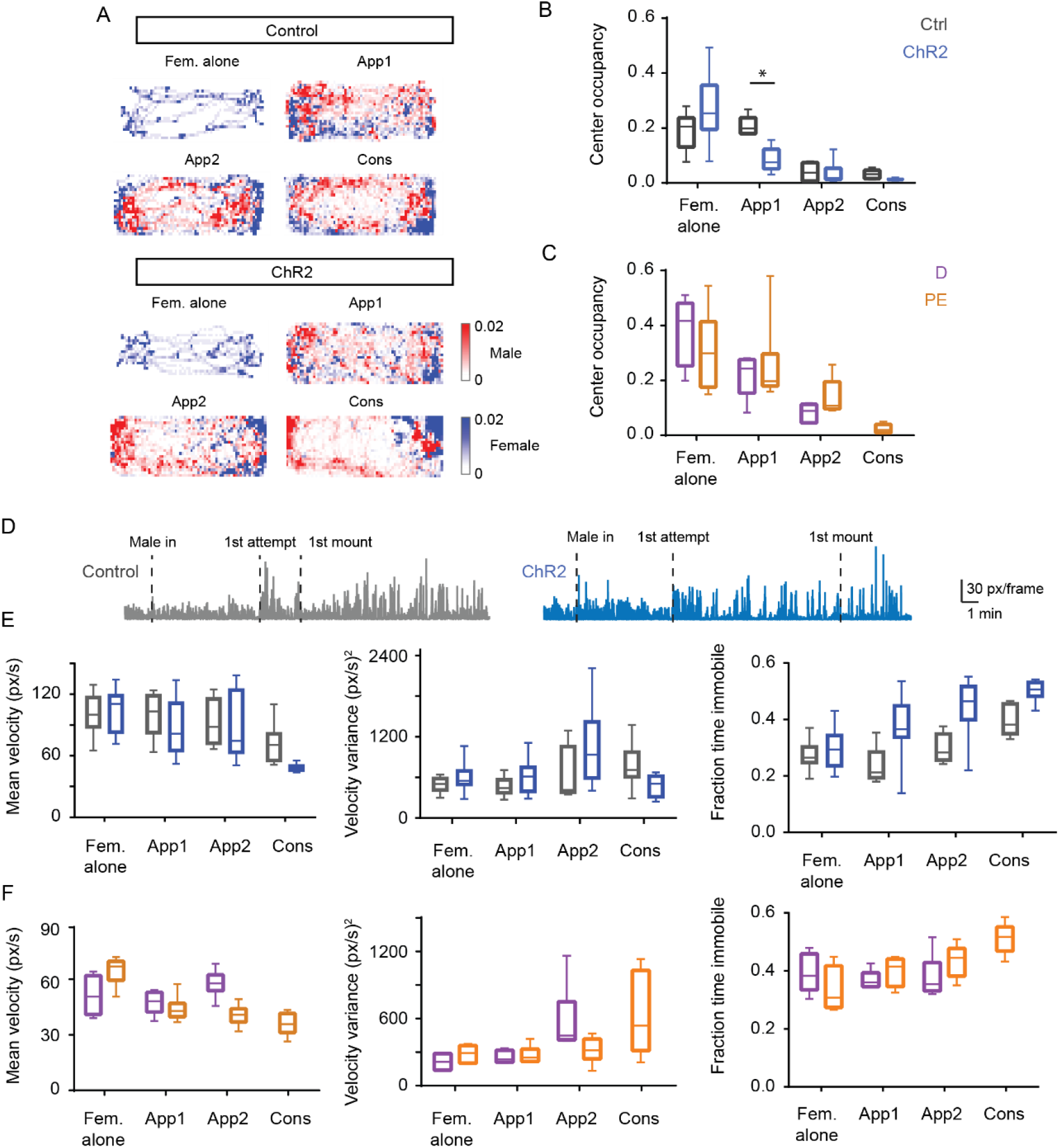
Cage occupancy and velocity are largely similar between ChR2, control, D and PE females. (A) Binned female and male occupancies throughout the different epochs of an example ChR2 and Ctrl session (B,C) Fraction of total time spent in the cage center throughout the different epochs in the optogenetics experiment (Ctrl vs. ChR2 females) and in the photometry experiment (D vs. PE females). * represents p<0.05 using Sidak’s multiple comparisons test. (D) Female velocity throughout an example Ctrl (left) and ChR2 (right) session. (E,F) Mean (left) and variance of the velocity (center) and the fraction of time immobile in the optogenetics experiment (E: Ctrl vs. ChR2 females) and in the photometry experiment (F: D vs. PE females).

## Methods

### Animals and reproductive/estrous cycle monitoring

Data were collected from adult (2–9 months) C57BL/6 (JAX stock #000664) (for cFos experiment and OVX vs NC-D socio-sexual assay), B6129S(Cg)-Pgrtm1.1(Cre)Shah/AndJ75 (PR-Cre^11^; JAX stock #017915) (photometry and optogenetics experiments), expressing Cre recombinase under the control of the progesterone receptor (PR) gene or PR-Cre-R26R-EYFP (for short PR-EYFP) (for electrophysiology experiments) female mice expressing the Enhanced Yellow Fluorescent Protein (EYFP) in cells expressing PR using the Cre-lox recombination system. Briefly, PR-EYFP mice result from the cross-breeding of PR-Cre mice with B6.129X1-Gt(ROSA26Sortm1(EYFP)Cos/J177 mice (for short R26R-EYFP; JAX stock #006148) that have the EYFP gene following a STOP sequence flanked by loxP.

Animals were kept under controlled temperature of 23±1 °C, reversed photoperiod of 12 h light/dark cycle (light available from 8 pm to 8 am) and group-housed conditions (unless specified otherwise) in standard cages with environmental enrichment elements. Food and water were provided *ad libitum*. Females were weaned at 20–21 days of age and group-housed with two to five animals. After reaching 6 weeks of age, females were exposed to adult C57BL/6 male soiled bedding once per week to stimulate the natural reproductive cycle. Cages were not changed on experimental days.

Females were habituated to pap smear collection daily for at least 4–5 days (approx. length of 1 complete reproductive cycle) before the experimental day. Pap smear collection consisted of manual restraint of the female followed by a lavage of the vaginal region (10 μL 0.1 mM PBS) which was then collected onto a slide. These smears were then stained using the Papanicolaou staining protocol^50^, and the reproductive state was determined by manual assessment of the proportion of different cell types in the stained smear^5^. Females were classified as D/diestrus or non-receptive when the predominant cell type was leukocytes, and as between PE/proestrus and estrus or receptive when there was a higher number of a-nucleated cornified cells along with an equal or lower proportion of nucleated epithelial cells in the smear. Experiments were performed immediately after pap smear collection and staining if the female was assessed as being in the PE state. Females that did not exhibit proper reproductive cycles during the habituation phase were not included in the study.

C57BL/6J male mice that had at least 3 prior ejaculation experiences within 3–4 weeks were used as studs for sexual behavior. For urine preference tests, urine was collected from age matched C57BL/6J males and females that would not be used for any other purpose during experiments.

Procedures were executed in accordance with the standards approved by the Commission for Experimentation and Animal Welfare of the Champalimaud Centre for the Unknown (Órgão para o Bem Estar Animal; ORBEA) and by the Portuguese National Authority for Animal Health (Direcção Geral de Alimentação e Veterinária; DGAV) (Ref. 0421/000/000/2018).

### Behavioral assay and cFos detection

For cFos detection, Adult C57BL/6 mice (N=23), comprising 18 sexually naïve test females, around 2 to 4 months old (PE: N=9; D: N=9), and 5 unrelated stud males, were used in a social/sexual paradigm.

Two females were tested per day within the 12:00 to 16:30 period: a PE, followed by a D. Females were allowed to habituate to the test cage for 10 minutes before the stud male was introduced. PE females interacted and mated with the stud for 5 mounts with intromissions or until ejaculation (if reached before), while D females interacted with the male for the same amount of time as the PE female or until 5 rejection events (including kicking and punching).

After 90 minutes, females were deeply anesthetized with sodium pentobarbital (120 mg/kg of body weight) and perfused transcardially with 0.1M PBS and 4% paraformaldehyde in 0.1M PBS. Brains were manually dissected, post-fixed in the same perfusion solution at 4°C overnight and later transferred into a 30% sucrose cryoprotectant solution.

Brains were cut into 45-μm-thick coronal sections using a microtome (SM 2000 R, Leica). Immunostaining was performed using cFos expression as a marker for neural activity in the VMH during the behavior assay. Floating VMH sections were incubated with primary cFos antibody (1:4000; 226 003, Synaptic Systems), followed by secondary antibody Alexa Fluor 488 Goat anti-Rabbit (1:1000; Abcam ab150077) and mounting in Mowiol (Sigma).

Sections were imaged using an AxioImager M2 (Zeiss) microscope and processed with ZEN 3.0 software (Zeiss). The location of brain regions was manually determined using the Allen Brain Atlas as a reference in Adobe Photoshop CC version 19.0 (Adobe). cFos positive cells were manually counted per VMHvl and VMHdm subregion (anterior, medial and posterior) and normalized to the area, calculated in ImageJ 1.52n.

### Behavioral assay with non-primed ovariectomized and diestrus naturally cycling females

Adult C57BL/6 mice were used in this behavioral assay: non-primed ovariectomized (OVX) females: N=3; naturally cycling D (NC-D) females: N=3, and 6 unrelated vasectomized stud males. We ovariectomized 2–3 months old C57BL/6 mice and, after their recovery, they were hormonally primed with estrogen (1 mg/ml) once per week until the week before the experiment (females were behaviorally tested 10–11 days after the last injection of estrogen). OVX females underwent the same handling and vaginal lavage as the NC females. Females were paired tested (NC-D and OVX within the same day) during the 12:00 and 17:30h period. Females were allowed to habituate to the test cage for 5 minutes before the stud male was introduced. The females and stud males interacted for 20 minutes after the first mount attempt.

### Behavioral annotation

Behaviors during the sexual behavior assays were manually annotated frame-by-frame using Python Video Annotator (Software Platform, Champalimaud Research). The following behavioral events were scored:

1. Male entry - point event - when the male was added to the cage and all 4 of his paws hit the cage floor
2. Anogenital investigation - point event - when one animal’s nose is in the vicinity of the anogenital region of the other animal. This could be a male to female anogenital investigation event or vice-versa
3. Mount attempt - point event - when the male puts his paws on the back of the female in order to mount her, but fails to perform shallow thrusts (probing) or thrusting
4. Mount without thrust - window of time - starts when the male puts his paws on the back of the female for a mount attempt but subsequently also performs probing, but not thrusts. Ends when the male removes both paws from the female’s back
5. Mount with thrust - window of time - starts when the male puts his paws on the back of the female, same as a mount attempt, and is able to perform thrusts. Ends when the male removes his paws from the female’s back
6. Thrust - point event - when the male’s body is most elongated once he is retreating from a thrust
7. Ejaculation - point event - when the male enters into the last thrust before shivering and falling to the side
8. Fight - point event - when the female lifts her hind- or fore-paw to hit the male
9. Escape - point event - when the female begins to run away from a mount attempt of a male and is successful in doing so within 15 frames of the mount attempt

The distance between the two animals was calculated using frame-by-frame 2-body centroid tracking in Bonsai, followed by analysis using a Python code.

### *In vivo* fiber photometry imaging

For photometry recordings, blue light from a fiber-coupled LED light source (465 nm, LEDFLP-465-560 FC, Doric Lenses) was bandpass filtered (passing band: 465±30 nm, FWHM-D25mm) and used to excite GCaMP6s. A set of three dichroic mirrors (Di01-R561-25×36; FF552-56 Di02-25×36; T495LP) allowed for light delivery and emitted fluorescence detection. Emitted fluorescence was bandpass filtered (passing band: 506±534 nm, FF02-520/28-25), reflected by the dichroic mirrors and detected by a photodetector (Doric). Photodetector signal was converted using a digital acquisition board (NI USB-6212, National Instruments) and recorded using Bonsai^51^ at a 1 kHz sampling rate.

Females were unilaterally injected with 300 nL of AA1-hSyn-Flex-GCaMP6s-WPre-SV40 (Addgene, #100845) in the aVMHvl (AP: −1.10, ML: –0.5, DV: −5.60; N=6) or in the pVMHvl (AP: −1.5, ML: –0.75, DV: −5.60; N=4) at 27.6 nL/min using a nanoliter injector (Nanoject II, Drummond Scientific) at 0.1 Hz. Subsequently, a 200 μm diameter mono fiber-optic cannula (MFC_200/245-0.37_8mm_ZF1.25_FLT, Doric Lenses) was unilaterally implanted 150–200 μm above the virus injection site and fixed using the light-cured dental cement Optibond (Optibond Universal, Kerr Dental, 36519) and Tetric EvoFlow (Tetric Evoflow Universal, Ivoclar Vivadent, 595953). 300 μL i.p. of saline and 100 μL s.c. of buprenorphine was administered 0.5 hours before the end of each surgery, and cages were left on a heating pad for 24 hours. Females were thereafter singly housed and allowed to recover for at least 2 weeks before pap smear habituation was initiated.

Female subjects underwent four experimental sessions: two in the D, and two in the PE state. In the current study we discuss the last two sessions, where the first session was D and the second session PE. Experiments were conducted in an experimental arena the same size as the home cage. Video recording was performed using a Point Grey camera (Flea3, Monochrome Point Grey) at 30 frames/second. A custom-built data acquisition and synchronization board was used for triggering the LED and video cameras and was controlled using a custom program written in Bonsai 2.4.0 51.

Each experimental session began with 5 minutes of habituation to the experimental arena, followed by 5 minutes of exposure to an object (a purple nitrile glove folded into a rectangular shape) and then 5 minutes of exposure to a female stimulus. After this the male was introduced into the arena for the sexual behavior assay. A 1 minute interval was given between each stimulus exposure. The D session was terminated 10–15 minutes after the first mount attempt by the male. The PE session was terminated either right after the male ejaculated, or 60 minutes after the first mount attempt by the male, whichever was the earlier time point.

Fluorescence traces were first corrected for bleaching over time by performing a rolling ΔF/F operation using the mean value of the lowest 20th percentile values within a 5-minute rolling window centered around each time point as the Fzero. For the first and last 2.5 minutes of the session, the Fzero used was that of the closest value calculable. This corrected GCaMP6s fluorescence trace was then used for further calculations. For the peri-event time histograms (PETHs), the relative change in GCaMP6s response (Z score) was calculated using the formula (F−mean(Fzero))/std(Fzero), where Fzero is the series of fluorescence values of GCaMP6s during a 2-second window prior to behavioral event onset. Peak Z score was calculated as the peak of the Z scored PETH within 3 seconds after behavioral onset. In Fig. 1G, a rolling-window Z score operation was performed on raw GCaMP6s traces, using the fluorescence values within a 5-minute rolling window centered around each time point as the Fzero and the same formula as for the PETHs. For plotting, all traces were smoothed using a 4th order Butterworth filter.

### *Ex vivo* electrophysiological recordings

After pap smearing and reproductive cycle determination, adult PR-Cre-EYFP female mice were decapitated under deep isoflurane anesthesia. The brains were quickly removed and placed into “ice cold” slicing solution containing (in mM): 0.66 kynurenic acid, 3.63 pyruvate, 2.5 KCl, 1.25 NaH2PO4, 26 NaHCO3, 10 D-Glucose, 230 Sucrose, 0.5 CaCl2, 10 MgSO4, and bubbled with 5% CO2 and 95% O2. The front third of the brain and the cerebellum were removed and coronal sections with 300μm containing the VMH were cut using a vibratome (Leica VT1200) in the same slicing solution. Slices were recovered in oxygenated artificial cerebrospinal fluid (ACSF) containing (in mM): 127 NaCl, 2.5 KCl, 25 NaHCO3, 1.25 NaH2PO4, 25 D-Glucose, 2 CaCl2, and 1 MgCl2, at 34°C for 30 minutes and stored in the same solution at room temperature before start recording.

Whole-cell recordings were made under a SliceScope Pro (Scientifica) microscope with the slices submerged in circulating ACSF. Patch recording pipettes (resistance 3-5 MΩ) were filled with internal solution containing (in mM) either: 135 K-Gluconate, 10 HEPES, 10 Na-phosphocreatine, 3 Na-L-ascorbate, 4 Mg-Cl2, 4 Na-ATP, 0.4 Na-GTP or 115 Cs-Methanesulfonate, 20 Cs-Cl,10 HEPES, 10 Na-phosphocreatine, 4 Na-ATP, 0.4 Na-GTP, 2.5 Mg-Cl2 and 0.6 EGTA (pH 7.2 adjusted with NaOH and osmolarity ∼292mOs) and 0.1% of biocytin. Internal solution was filtered with a 0.2μm pore size cellulose acetate filter tip. A Multiclamp 700B amplifier and digitized at 10KHz with a Digidata 1440a digitizer (both from Molecular Devices) were used and the data was filtered online with a 5 KHz low-pass filter.

We applied a test-pulse of –10 mV for 100 ms in voltage-clamp mode at –70 mV to monitor membrane and series resistance. Excitatory postsynaptic currents (EPSCs) were recorded in voltage-clamp mode at –70mV holding potential (GABA-receptors Chloride reversal potential, as calculated using Nernst equation) and spontaneous inhibitory postsynaptic currents (IPSCs) at 0mV holding potential (AMPA receptors reversal potential). Spontaneous and miniature synaptic events were recorded during 2–3 minutes. Electrically evoked synaptic currents were evoked by delivering 0.5 ms pulses of current in the vicinity of the recorded neuron through a glass micropipette (3–5 MΩ) connected to a microstimulation device (ISO-Flex, AMPI). Stimulation intensity was manually adjusted to evoke EPSC responses of around 100–300 pA of amplitude while avoiding escaped spikes in voltage clamp mode. To assess paired pulse facilitation, 2 consecutive pulses were delivered at 25 Hz, followed by an interval of 8 seconds before the following paired stimulation.

To functionally map GABAergic synapses, ACSF containing 150 µM RuBi-GABA trimethylphosphine (Tocris) was circulated in the slice recording chamber. Using galvanometer mirrors (LASU system, Scientifica), we quickly positioned a blue laser beam (LASU system, Scientifica) switching over a grid of 14×14 locations (80 µm apart) at 1 Hz in semi-randomized order to maximize the distance between consecutive stimulation locations. The laser was delivered through an air-immersed objective (Olympus, 4x, NA 0.1) at a fixed power (8 mW at the objective tip). The resulting data per cell was the average of 3 repetitions of the 14×14 grid.

### Electrophysiology data analysis

Voltage clamp test-pulses were analyzed using Clampfit 10.7 Software (Molecular Devices, LLC). Access resistance (Ra) was determined by measuring the amplitude of the fast current response to the command voltage step and membrane resistance (Rm) as the difference between the baseline and the holding current in the steady state after the capacitive decay, by applying Ohm’s law. Only neurons with Ra 1/10th lower than the Rm were used for analysis. Ra was comparable across location, genotype and the phase of the cycle in the present study.

Electrophysiological recordings of spontaneous and miniature EPSCs and IPSCs were analyzed using the template search function of Clampfit 10.7 Software (Molecular Devices, LLC.). The amplitude of evoked currents was analyzed using Clampfit 10.7 Software (Molecular Devices, LLC.) after manually removing sweeps in which escaped spikes were observed. Both for amplitudes and frequencies, the excitation to inhibition balance index was calculated using the following formula: EI index= (E–I) / (E+I)

Uncaging recordings were imported into MATLAB and analyzed using custom written software. A minimum of 3 uncaging maps were averaged per neuron, the summed input of active pixels (locations in which the peak amplitude was larger than 4 times the standard deviation of the baseline noise amplitude) was obtained for those pixels closer than (proximal) or larger than (distal) 250 µm from the soma in cartesian coordinates.

### Neuronal reconstruction and image data analysis

32 out of 46 VMHvl neurons recorded using uncaging of GABA were filled with biocytin in acute slices obtained from PR-EYFP female mice using whole-cell recording pipettes. These slices containing biocytin filled neurons were fixed and stained with streptavidin-Alexa Fluor 594 (1:1000, 2h incubation at RT). We used a previously described protocol to analyze the morphologic properties of the neurons filled with biocytin^18^. Briefly, we acquired confocal stack images of the biocytin labeled neurons (Zeiss LSM 710 confocal laser scanning microscope) and used the simple neurite tracer^52^ package from Fiji/ImageJ software to analyze their morphologic properties.

### *In vivo* optogenetic manipulations

For aVMHvlPr+ neuron stimulation, females were bilaterally injected with 75 nL of AAV2.EF1a.Flex.ChR3.EYFP (UNC vector core) for the ChR2 group (N=11) or AAV2.EF1a.Flex.EYFP (UNC vector core) for the Ctrl group (N=6) in the aVMHvl (AP: −1.10, ML: ±0.5, DV: −5.50) at 27.6 nL/min using a nanoliter injector (Nanoject II, Drummond Scientific) at 0.1 Hz. Subsequently, 200 μ diameter homemade cannulas were bilaterally implanted 150 μm above the virus injection site. Implanted cannulas were bent outwards/laterally to allow space for both cannulas and were fixed using the light-cured dental cement Optibond (Optibond Universal, Kerr Dental, 36519) and Tetric EvoFlow (Tetric Evoflow Universal, Ivoclar Vivadent, 595953). 300 μL i.p. of saline and 100 μL s.c. of buprenorphine was administered 0.5 hours before the end of each surgery, and cages were left on a heating pad for 24 hours. Females were thereafter singly housed and allowed to recover for at least 2 weeks before pap smear habituation was initiated.

Light was delivered to the cannulas using a 450 nm LED (LDFLS_450/075, Doric Lenses) via a 200 μ splitter branching patch cord (Doric Lenses). Light stimulation (20 ms, 20 Hz, 5 mW) was delivered for 30 seconds every minute (30 sec on – 30 sec off) throughout the session. Before each session, the LED power during pulsing was measured at each tip of the patch cord using an optical power meter (PM130D, Thorlabs, Inc.) and set to 5 mW.

All experiments were performed on PE females. The experimental session began with 5 minutes in the home cage, followed by an open field test (OFT). In the OFT, the female was introduced into one corner of the open field arena (dimensions: 40 cm X 40 cm X 25 cm) and allowed to explore freely for 5 minutes. After this, a 5-minute male/female urine preference test was conducted, where 2 small petri dishes (60 mm diameter, Greiner Bio-One) were introduced into the central portion of the arena, each of which had filter papers soaked in 20 μL of male or female urine. The urine spots had a mixture of urine from 2 males/females that were unrelated to any other aspect of the experimental paradigm. Following this, the female was transferred to the experimental arena, and after a minute, a male stud was added into the arena. The session was terminated either right after the male ejaculated, or 30 minutes after the first mount attempt by the male if the female did not allow the male to successfully mount her until then; we based our decision to interrupt the session 30 min after the first copulation attempt. All our males were screened for sexual performance and we based the interval on previous studies of the lab ^53, 54^ where we observed that after the first mount attempt, all males ejaculated in less than 10 minutes. In the latter case a follow up pap smear was collected immediately after the conclusion of the experiment to assess if the female had transitioned to the next phase of the reproductive cycle, which would explain her non-receptive behavior. Only females that had follow-up pap smears still in the PE state were included. If the male did not exhibit a mount attempt within 30 minutes of introduction into the cage, he was removed and another male was added to the cage.

Video recording was performed using 2 cameras (Flea3, Monochrome, Point Grey), one top view and one side view, at 30 frames/second. A custom data acquisition and synchronization board was used for triggering the LED and video cameras and was controlled using a custom program written in Bonsai 2.4.0.

On the day of sacrifice, females underwent 10 minutes of the experimental light stimulation protocol in the experimental cage before being returned to their home cage. Perfusion was performed after 90 minutes using PBS followed by fixation using ice-cold 0.4% PFA. Heads were severed and stored whole at 4℃ in 0.4% PFA for 24–48 h before the brains were removed and stored in 30% sucrose-PBS + 0.1% sodium azide solution until being sliced.

Brains were coronally sliced on a sliding microtome (SM2000 R, Leica) or a cryostat (CM1800, Leica) at 45–50 μ thickness. Brain slices then underwent a double immunostaining for EYFP and cFos using a primary antibody cocktail of rabbit anti-cFos (1:2000, Synaptic Systems, Cat. No. 226 008) and goat anti-GFP (1:1000, Abcam, ab6673) followed by a secondary antibody cocktail of Alexa Fluor 594 donkey anti-rabbit (1:1000, Invitrogen, ab150076) and Alexa Fluor Plus 488 donkey anti-goat (1:1000, Invitrogen, A32814). Slices were mounted on glass slides (Thermo Fisher Scientific) using mowiol (Sigma-Aldrich) as a mounting medium, cover-slipped (Thermo Scientific Menzel) and kept at 4℃ until imaging.

To determine virus and fiber location, images were acquired using a wide-field motorized microscope (Axio Imager 2, Zeiss). For quantification of ChR2/EYFP and cFos co-expression, images were acquired using a Zeiss LSM 710 confocal laser scanning microscope at 20x magnification. Brain slices containing the aVMHvl were identified by overlaying the Allen brain atlas on brain slice images in Adobe Photoshop (Adobe). A representative brain slice from each animal was used for the quantification of ChR2/EYFP and cFos expression. Cell counting was performed manually using ImageJ.

The position of the female in behavioral videos was tracked using custom Bonsai programs. For OFT analysis, a region of interest (ROI) corresponding to the central region of the open field which had area equal to half the area of the entire box was delineated and every frame in which the animal entered this ROI was recorded. The centroid position of the female was used to calculate the distance traveled by the female using a Python script. For urine preference test analysis, ROIs were drawn around the petri dishes. Every frame where the female entered the ROIs was recorded and this was used to calculate the percentage of time spent interacting with male vs. female urine.

For behavioral session subsampling analysis, we first calculated the median duration of appetitive 2 in Ctrl females as 30 seconds. We used two times this duration, i.e., 60 seconds as a threshold and every animal that had an appetitive 2 duration greater than this threshold underwent subsampling. For these animals we chose 100 random 30-second samples from the appetitive 2 phase and used the mean of the behavioral parameter during these 30-second samples as the mean of appetitive 2 for that animal. The rest of the analysis remained the same.

### Markerless pose tracking

We used SLEAP^35^ for tracking the identity and position of the male and female during the socio-sexual interaction assays. We trained a bottom up model using a UNET backbone. The training set consisted of 230 annotated frames throughout 12 videos in which, for both male and female, the following body parts were indicated: nose, implant/ head base, right ear, left ear, back 1, back 2, tail base, tail 1, tail 2, right shoulder, right hip, left shoulder, left hip.

The frames for the training set were selected using the image feature method. The default training parameters were kept except the batch size, which was set to 12 and the rotation angle of the augmentation phase, that was set to [–180 180]. After training the model, the inference was done setting to 2 the number of instances per frame and using optical flow for tracking the cross-frame identities (through keypoint similarity using the Hungarian matching method). Predicted frames were visually inspected and identity switches were manually corrected when present. Predictions were then exported to H5 files and loaded into MATLAB for analysis using custom written software.

### Statistical analysis

Statistical analyses were performed using MATLAB, Python or GraphPad Prism 9 Software. Normality of the residuals was tested with the Shapiro-Wilk test. In data that passed the normality test, independent-samples Student’s t-test were used to evaluate differences between groups. If data did not follow a normal distribution, analysis was performed using a non-parametric Mann– Whitney U test for unpaired samples.

Box plots indicate median and the interquartile range (IQR, ±25th–75th percentile) and the whisker edges represent the minimum and maximum data limits excluding outliers using the Tukey criterion (outliers were depicted outside the box plot). Error bars and shaded error bars represent mean and standard error of the mean. For comparing fractions (i.e. percentage of animals that had sex or fought), we used the Z test. Paired measurements were analyzed with a paired t-test. Two-way ANOVA tests were performed to compare groups in different phases of the cycle (D vs. PE) versus location in the AP axis (anterior vs. medial vs. posterior), using the Sidak test to correct for multiple comparisons. Statistical significance was set at p<0.05.

## References

1. Dey, S., Chamero, P., Pru, J.K., Chien, M.-S., Ibarra-Soria, X., Spencer, K.R., Logan, D.W., Matsunami, H., Peluso, J.J., and Stowers, L. (2015). Cyclic Regulation of Sensory Perception by a Female Hormone Alters Behavior. Cell 161, 1334–1344. 10.1016/j.cell.2015.04.052.

2. Inoue, S., Yang, R., Tantry, A., Davis, C., Yang, T., Knoedler, J.R., Wei, Y., Adams, E.L., Thombare, S., Golf, S.R., et al. (2019). Periodic Remodeling in a Neural Circuit Governs Timing of Female Sexual Behavior. Cell 179, 1393–1408.e16. 10.1016/j.cell.2019.10.025.

3. Pfaff, D.W., Kow, L.-M., Loose, M.D., and Flanagan-Cato, L.M. (2008). Reverse engineering the lordosis behavior circuit. Horm. Behav. 54, 347–354. 10.1016/j.yhbeh.2008.03.012.

4. Mong, J.A., and Pfaff, D.W. (2004). Hormonal symphony: steroid orchestration of gene modules for sociosexual behaviors. Mol. Psychiatry 9, 550–556. 10.1038/sj.mp.4001493.

5. Gutierrez-Castellanos, N., Husain, B.F.A., Dias, I.C., and Lima, S.Q. (2022). Neural and behavioral plasticity across the female reproductive cycle. Trends Endocrinol. Metab. 0. 10.1016/j.tem.2022.09.001.

6. Lenschow, C., and Lima, S.Q. (2020). In the mood for sex: neural circuits for reproduction. Curr. Opin. Neurobiol. 60, 155–168. 10.1016/j.conb.2019.12.001.

7. Pfaff, D.W., and Sakuma, Y. Deficit in the lordosis reflex of female rats caused by lesions in the ventromedial nucleus of the hypothalamus. 8.

8. Pfaff, D.W., and Sakuma, Y. FACILITATION OF THE LORDOSIS REFLEX OF FEMALE RATS FROM THE VENTROMEDIAL NUCLEUS OF THE HYPOTHALAMUS. 14.

9. Bentefour, Y., and Bakker, J. (2021). Kisspeptin signaling and nNOS neurons in the VMHvl modulate lordosis behavior but not mate preference in female mice. Neuropharmacology 198, 108762. 10.1016/j.neuropharm.2021.108762.

10. McCarthy, M.M., Malik, K.F., and Feder, H.H. (1990). Increased GABAergic transmission in medial hypothalamus facilitates lordosis but has the opposite effect in preoptic area. Brain Res. 507, 40–44. 10.1016/0006-8993(90)90519-H.

11. Yang, C.F., Chiang, M.C., Gray, D.C., Prabhakaran, M., Alvarado, M., Juntti, S.A., Unger, E.K., Wells, J.A., and Shah, N.M. (2013). Sexually dimorphic neurons in the ventromedial hypothalamus govern mating in both sexes and aggression in males. Cell 153, 896–909. 10.1016/j.cell.2013.04.017.

12. Hashikawa, K., Hashikawa, Y., Tremblay, R., Zhang, J., Feng, J.E., Sabol, A., Piper, W.T., Lee, H., Rudy, B., and Lin, D. (2017). Esr1+ cells in the ventromedial hypothalamus control female aggression. Nat. Neurosci. 20, 1580–1590. 10.1038/nn.4644.

13. Yin, L., Hashikawa, K., Hashikawa, Y., Osakada, T., Lischinsky, J.E., Diaz, V., and Lin, D. (2022). VMHvllCckar cells dynamically control female sexual behaviors over the reproductive cycle. Neuron 0. 10.1016/j.neuron.2022.06.026.

14. Nomoto, K., and Lima, S.Q. (2015). Enhanced Male-Evoked Responses in the Ventromedial Hypothalamus of Sexually Receptive Female Mice. Curr. Biol. 25, 589–594. 10.1016/j.cub.2014.12.048.

15. Lenschow, C., Mendes, A.R.P., and Lima, S.Q. (2022). Hearing, touching, and multisensory integration during mate choice. Front. Neural Circuits 16.

16. Elias, L.J., Schaffler, M., Succi, I., Foster, W., Gradwell, M., Bohic, M., Ejoh, L., Abraira, V., and Abdus-Saboor, I. (2022). Identification of touch neurons underlying dopaminergic pleasurable touch and sexual receptivity. 2021.09.22.461355. 10.1101/2021.09.22.461355.

17. Rubin, B.S., and Barfield, R.J. (1983). Progesterone in the Ventromedial Hypothalamus Facilitates Estrous Behavior in Ovariectomized, Estrogen-Primed Rats*. Endocrinology 113, 797–804. 10.1210/endo-113-2-797.

18. Dias, I.C., Gutierrez-Castellanos, N., Ferreira, L., and Lima, S.Q. (2021). The Structural and Electrophysiological Properties of Progesterone Receptor-Expressing Neurons Vary along the Anterior-Posterior Axis of the Ventromedial Hypothalamus and Undergo Local Changes across the Reproductive Cycle. eNeuro 8, ENEURO.0049-21.2021. 10.1523/ENEURO.0049-21.2021.

19. Calizo, L.H., and Flanagan-Cato, L.M. (2002). Estrogen-induced dendritic spine elimination on female rat ventromedial hypothalamic neurons that project to the periaqueductal gray. J. Comp. Neurol. 447, 234–248. 10.1002/cne.10223.

20. Ferri, S.L., Hildebrand, P.F., Way, S.E., and Flanagan-Cato, L.M. (2014). Estradiol regulates markers of synaptic plasticity in the hypothalamic ventromedial nucleus and amygdala of female rats. Horm. Behav. 66, 409–420. 10.1016/j.yhbeh.2014.06.016.

21. Booth, C., Wayman, C.P., and Jackson, V.M. (2010). An ex vivo multi-electrode approach to evaluate endogenous hormones and receptor subtype pharmacology on evoked and spontaneous neuronal activity within the ventromedial hypothalamus; translation from female receptivity. J. Sex. Med. 7, 2411–2423. 10.1111/j.1743-6109.2010.01843.x.

22. Lo, L., Yao, S., Kim, D.-W., Cetin, A., Harris, J., Zeng, H., Anderson, D.J., and Weissbourd, B. (2019). Connectional architecture of a mouse hypothalamic circuit node controlling social behavior. Proc. Natl. Acad. Sci. 116, 7503–7512. 10.1073/pnas.1817503116.

23. Kim, D.-W., Yao, Z., Graybuck, L.T., Kim, T.K., Nguyen, T.N., Smith, K.A., Fong, O., Yi, L., Koulena, N., Pierson, N., et al. (2019). Multimodal Analysis of Cell Types in a Hypothalamic Node Controlling Social Behavior. Cell 179, 713–728.e17. 10.1016/j.cell.2019.09.020.

24. Wang, L., Talwar, V., Osakada, T., Kuang, A., Guo, Z., Yamaguchi, T., and Lin, D. (2019). Hypothalamic Control of Conspecific Self-Defense. Cell Rep. 26, 1747–1758.e5. 10.1016/j.celrep.2019.01.078.

25. Liu, M., Kim, D.-W., Zeng, H., and Anderson, D.J. (2022). Make war not love: The neural substrate underlying a state-dependent switch in female social behavior. Neuron 110, 841–856.e6. 10.1016/j.neuron.2021.12.002.

26. Blanchard, R.J., Hebert, M.A., Ferrari, P.F., Palanza, P., Figueira, R., Blanchard, D.C., and Parmigiani, S. (1998). Defensive behaviors in wild and laboratory (Swiss) mice: the mouse defense test battery. Physiol. Behav. 65, 201–209. 10.1016/s0031-9384(98)00012-2.

27. Zinck, L., and Lima, S.Q. (2013). Mate Choice in Mus musculus Is Relative and Dependent on the Estrous State. PLOS ONE 8, e66064. 10.1371/journal.pone.0066064.

28. Pfaus, J.G. (1996). Frank A. Beach award. Homologies of animal and human sexual behaviors. Horm. Behav. 30, 187–200. 10.1006/hbeh.1996.0024.

29. Ball, G.F., and Balthazart, J. (2008). How useful is the appetitive and consummatory distinction for our understanding of the neuroendocrine control of sexual behavior? Horm. Behav. 53, 307–318. 10.1016/j.yhbeh.2007.09.023.

30. Bullitt, E. (1990). Expression of c-fos-like protein as a marker for neuronal activity following noxious stimulation in the rat. J. Comp. Neurol. 296, 517–530. 10.1002/cne.902960402.

31. Mhaouty-Kodja, S., Naulé, L., and Capela, D. (2018). Sexual Behavior: From Hormonal Regulation to Endocrine Disruption. Neuroendocrinology 107, 400–416. 10.1159/000494558.

32. Rubin, B.S., and Barfield, R.J. (1983). Induction of estrous behavior in ovariectomized rats by sequential replacement of estrogen and progesterone to the ventromedial hypothalamus. Neuroendocrinology 37, 218–224. 10.1159/000123546.

33. Glasgow, S.D., McPhedrain, R., Madranges, J.F., Kennedy, T.E., and Ruthazer, E.S. (2019). Approaches and Limitations in the Investigation of Synaptic Transmission and Plasticity. Front. Synaptic Neurosci. 11.

34. Rial Verde, E.M., Zayat, L., Etchenique, R., and Yuste, R. (2008). Photorelease of GABA with Visible Light Using an Inorganic Caging Group. Front. Neural Circuits 2, 2. 10.3389/neuro.04.002.2008.

35. Pereira, T.D., Tabris, N., Matsliah, A., Turner, D.M., Li, J., Ravindranath, S., Papadoyannis, E.S., Normand, E., Deutsch, D.S., Wang, Z.Y., et al. (2022). SLEAP: A deep learning system for multi-animal pose tracking. Nat. Methods 19, 486–495. 10.1038/s41592-022-01426-1.

36. Gao, Z., van Beugen, B.J., and De Zeeuw, C.I. (2012). Distributed synergistic plasticity and cerebellar learning. Nat. Rev. Neurosci. 13, 619–635. 10.1038/nrn3312.

37. Knoedler, J.R., Inoue, S., Bayless, D.W., Yang, T., Tantry, A., Davis, C., Leung, N.Y., Parthasarathy, S., Wang, G., Alvarado, M., et al. (2022). A functional cellular framework for sex and estrous cycle-dependent gene expression and behavior. Cell 185, 654–671.e22. 10.1016/j.cell.2021.12.031.

38. Stagkourakis, S., Spigolon, G., Liu, G., and Anderson, D.J. (2020). Experience-dependent plasticity in an innate social behavior is mediated by hypothalamic LTP. Proc. Natl. Acad. Sci. 117, 25789–25799. 10.1073/pnas.2011782117.

39. Krzywkowski, P., Penna, B., and Gross, C.T. (2019). Dynamic encoding of social threat and spatial context in the hypothalamus (Neuroscience) 10.1101/811380.

40. Arancio, O., Kiebler, M., Lee, C.J., Lev-Ram, V., Tsien, R.Y., Kandel, E.R., and Hawkins, R.D. (1996). Nitric Oxide Acts Directly in the Presynaptic Neuron to Produce Long-Term Potentiationin Cultured Hippocampal Neurons. Cell 87, 1025–1035. 10.1016/S0092-8674(00)81797-3.

41. Mani, S.K., Allen, J.M., Rettori, V., McCann, S.M., O’Malley, B.W., and Clark, J.H. (1994). Nitric oxide mediates sexual behavior in female rats. Proc. Natl. Acad. Sci. 91, 6468–6472. 10.1073/pnas.91.14.6468.

42. Donato, J., Jr., Frazão, R., Fukuda, M., Vianna, C.R., and Elias, C.F. (2010). Leptin Induces Phosphorylation of Neuronal Nitric Oxide Synthase in Defined Hypothalamic Neurons. Endocrinology 151, 5415–5427. 10.1210/en.2010-0651.

43. Faber, C.L., Matsen, M.E., Velasco, K.R., Damian, V., Phan, B.A., Adam, D., Therattil, A., Schwartz, M.W., and Morton, G.J. (2018). Distinct Neuronal Projections From the Hypothalamic Ventromedial Nucleus Mediate Glycemic and Behavioral Effects. Diabetes 67, 2518–2529. 10.2337/db18-0380.

44. Jo, Y.-H. (2012). Endogenous BDNF regulates inhibitory synaptic transmission in the ventromedial nucleus of the hypothalamus. J. Neurophysiol. 107, 42–49. 10.1152/jn.00353.2011.

45. Jamshidi, N., and Taylor, D.A. (2001). Anandamide administration into the ventromedial hypothalamus stimulates appetite in rats. Br. J. Pharmacol. 134, 1151–1154. 10.1038/sj.bjp.0704379.

46. Emmerson, P.J., and Miller, R.J. (1999). Pre- and postsynaptic actions of opioid and orphan opioid agonists in the rat arcuate nucleus and ventromedial hypothalamus in vitro. J. Physiol. 517, 431–445. 10.1111/j.1469-7793.1999.0431t.x.

47. Georgescu, M., and Pfaus, J.G. (2006). Role of glutamate receptors in the ventromedial hypothalamus in the regulation of female rat sexual behaviors: II. Behavioral effects of selective glutamate receptor antagonists AP-5, CNQX, and DNQX. Pharmacol. Biochem. Behav. 83, 333–341. 10.1016/j.pbb.2006.02.019.

48. Anderson, D.J. (2016). Circuit modules linking internal states and social behaviour in flies and mice. Nat. Rev. Neurosci. 17, 692–704. 10.1038/nrn.2016.125.

49. Tinbergen, N. The hierarchical organization of nervous mechanisms underlying instinctive behaviour. Symp. Soc. Exp. Biol. 4, 305–312 (1950).

50. Caligioni, C.S. (2009). Assessing reproductive status/stages in mice. Curr. Protoc. Neurosci. Appendix 4, Appendix 4I. 10.1002/0471142301.nsa04is48.

51. Lopes, G., Bonacchi, N., Frazão, J., Neto, J.P., Atallah, B.V., Soares, S., Moreira, L., Matias, S., Itskov, P.M., Correia, P.A., et al. (2015). Bonsai: an event-based framework for processing and controlling data streams. Front. Neuroinformatics 9.

52. Longair, M.H., Baker, D.A., and Armstrong, J.D. (2011). Simple Neurite Tracer: open source software for reconstruction, visualization and analysis of neuronal processes. Bioinforma. Oxf. Engl. 27, 2453–2454. 10.1093/bioinformatics/btr390.

53. Lenschow, C., Mendes, A.R.P., Ferreira, L., Lacoste, B., Quilgars, C., Bertrand, S.S., and Lima, S.Q. (2022). A galanin-positive population of lumbar spinal cord neurons modulates sexual behavior and arousal. 2022.10.04.510783. 10.1101/2022.10.04.510783.

54. Valente, S., Marques, T., and Lima, S.Q. (2021). No evidence for prolactin’s involvement in the post-ejaculatory refractory period. Commun. Biol. 4, 10. 10.1038/s42003-020-01570-4.

55 Yoshida, M. et al. (1994) Estrogen reduces the excitability of the female rat medial amygdala afferents from the medial preoptic area but not those from the lateral septum. Exp. Brain Res. 101, 1–7

